# Triptolide sensitizes cancer cells to nucleoside DNA methyltransferase inhibitors through inhibition of DCTPP1-mediated cell-intrinsic resistance

**DOI:** 10.1101/2024.05.19.594134

**Authors:** Jianyong Liu, Qing-Li He, Jianya Zhou, Roshan Chikarmane, Glenn Hauk, Archana Rachakonda, Ajay M. Vaghasia, Nicole Castagna, Ruchama C. Steinberg, Minh-Tam Pham, Nicole M. Anders, Teresia M. Wanjiku, Philipp Nuhn, Joong Sup Shim, Hugh Giovinazzo, David M. Esopi, Kunhwa Kim, Jonathan Coulter, Rulin Wang, Jianying Zhou, Michelle A. Rudek, James M. Berger, Jun O. Liu, William G. Nelson, Srinivasan Yegnasubramanian

**Affiliations:** The Sidney Kimmel Comprehensive Cancer Center at Johns Hopkins University, Baltimore, MD, 21231 USA; Department of Pharmacology and Molecular Sciences, School of Medicine, Johns Hopkins University, Baltimore, Maryland, 21231 USA; Department of Respiratory Disease, Thoracic Disease Center, The First Affiliated Hospital, College of Medicine, Zhejiang University, Hangzhou, China PR; Faculty of Health Sciences, University of Macau, Taipa, Macau SAR; Department of Medicine, Division of Clinical Pharmacology, School of Medicine, Johns Hopkins University, Baltimore, Maryland, 21231 USA; Institute for Basic Biomedical Sciences and Department of Biophysics, Johns Hopkins University School of Medicine, Baltimore, Maryland; inHealth Precision Medicine, Johns Hopkins Medicine, Baltimore, Maryland

## Abstract

While nucleoside DNA methyltransferase inhibitors (DNMTi) such as decitabine and azacitidine are effective in treating myelodysplatic syndrome (MDS)/leukemia, they have had limited utility for the majority of other cancers. Through a chemical library screen, we identified that triptolide, a diterpenoid epoxide from *Tripterygium wilfordii*, or analogs significantly augmented the epigenetic and anti-cancer effects of decitabine *in vitro* and *in vivo*. These effects were attributable to inhibition of DCTPP1-mediated cleavage of 5-aza-deoxycytidine triphosphate, the convergent activated metabolite of nucleoside DNMTi, leading to enhanced drug incorporation into genomic DNA, increased DNMT degradation, enhanced global DNA demethylation and associated transcriptional reprogramming. We show that high DCTPP1 expression was associated with cell-intrinsic resistance to nucleoside DNMTi, and that triptolide and its analogs could overcome this resistance.

**SIGNIFICANCE:** We screened a library of existing drugs to identify those capable of enhancing the anti-cancer effects of the nucleoside DNMTi decitabine. The combination of triptolide and decitabine synergistically inhibited cancer cell growth and survival *in vitro*, and was highly effective in inhibiting xenograft growth *in vivo*. Biochemical, genetic and structural biology studies with triptolide and its analogs revealed that this synergy was due to their inhibition of DCTPP1-mediated pyrophosphate cleavage from 5-aza-deoxycytidine triphosphate, the active metabolite of DNMTi. The genomic incorporation and efficacy of decitabine in cancer cell lines were significantly correlated with DCTPP1 expression more so than those of other nucleoside metabolizing genes. Triptolide and its analogs comprise rational adjuncts to nucleoside DNMTi ripe for further pre-clinical/clinical translation.

**HIGHLIGHTS:**

- Triptolide synergistically sensitizes cancer cells to DNMTi *in vitro*.
- Triptolide and decitabine combination shows favorable efficacy and safety *in vivo*.
- Synergy of triptolide and decitabine is mediated through inhibition of DCTPP1.
- High DCTPP1 expression confers cell intrinsic resistance to DNMTi.

## INTRODUCTION

In cancer cells, somatic alterations in DNA methylation appear to occur consistently, to arise early, and to be potentially reversible (Brooks et al., 1998; Herman and Baylin, 2003; Lin et al., 2001; Nakayama et al., 2003; Yegnasubramanian et al., 2004). Increased CpG dinucleotide methylation at transcriptional regulatory regions of many genes typically facilitates gene repression, leading to silencing of critical cancer genes (Antequera and Bird, 1993; Bird, 1986; Herman and Baylin, 2003). Suggestive of a driver role in human carcinogenesis and disease progression, such DNA hypermethylation alterations can be strongly maintained and subject to selection through disease progression and metastatic dissemination (Aryee et al., 2013; Chiappinelli et al., 2015; Yegnasubramanian et al., 2008; Yegnasubramanian et al., 2004). For this reason, therapeutic reversal of epigenetic gene silencing, with nucleoside analog DNA methyltransferase (DNMT) inhibitors such as decitabine (DAC, Dacogen®), azacitidine (Vidaza®), and newly developed oral formulations of these drugs (ASTX-727 (Inqovi®) and Onureg®), have been developed as rational anti-cancer treatment strategies. The mechanism of action of these drugs involves metabolic conversion to 5-aza-2’-deoxycytidine triphosphate(5-aza-dCTP), incorporation into genomic DNA, trapping of DNMT enzymes and their subsequent proteolytic degradation, and ultimately DNA demethylation through passive loss of 5-methylcytosine (Ghoshal et al., 2005; Patel et al., 2010; Yang et al., 2010). Thus far, the DNMTi have shown promise when used for myelodysplastic syndrome (MDS), for certain acute myeloid leukemias (AMLs), and a limited number of other cancers (Blum et al., 2007; Cashen et al., 2010; Issa et al., 2004; Kantarjian et al., 2006; Silverman and Mufti, 2005).

However, DNMTi have not been effective as single agents for the majority of cancer types despite the nearly universal prevalence of DNA hypermethylation alterations across cancer types (Cheishvili et al., 2015; Jones and Baylin, 2002). Additionally, the clinical use of nucleoside analog DNMTi has been generally limited by poor bioavailability, instability in blood, and substantial toxicity when administered at high doses (Karpf et al., 2001; Palii et al., 2008). The difficulties in administering higher nucleoside DNMTi doses has even obfuscated whether DNMT inhibition is responsible *per se* for the responses seen in hematologic disorders (Issa and Kantarjian, 2009). Some of the poor pharmacologic properties of the DNMTi have been attributed to rapid metabolism of the nucleoside analogs by cytidine deaminase (CDA) (Ebrahem et al., 2012; Qin et al., 2011; Zauri et al., 2015). Strategies aimed at inhibiting CDA in combination with decitabine or azacitidine to enhance the bioavailability and stability of the DNMTi have been translated to clinical use (Garcia-Manero et al., 2020; Issa et al., 2015; Lavelle et al., 2012; Lemaire et al., 2009; Savona et al., 2019). Lower doses of decitabine or azacitidine, given alone or in combination with histone deacetylase (HDAC) inhibitors, have exhibited improved safety with preserved efficacy in clinical trials (Chai et al., 2008; Kalac et al., 2011; Luszczek et al., 2010).

We hypothesized that there may be cancer cell intrinsic mechanisms that could confer primary or acquired resistance to DNA methyltransferase inhibitors, potentially limiting their utility in reversing DNA methylation and anti-cancer efficacy. To ascertain whether available drugs could inhibit such resistance mechanisms, we screened 3800 compounds from the Johns Hopkins Drug Library collection (Chong et al., 2006; Platz et al., 2011) alone or in combination with decitabine, and identified that triptolide (TPL), a diterpenoid epoxide found in the Thunder God Vine (*Tripterygium wilfordii*), exhibited marked synergy with decitabine in cancer cell growth inhibition and cytotoxicity. Using pharmacological/biochemical, molecular genetic, and biophysical approaches, we identified that inhibition of DCTPP1 by triptolide and its analogs accounted for the synergy between triptolide and decitabine. By inhibiting DCTPP1-mediated pyrophosphate cleavage from the active decitabine metabolite 5-aza-dCTP, triptolide/analogs facilitated greater drug incorporation into the genome, triggering increased DNMT degradation, enhanced demethylation and modulation of gene expression programs including interferon response pathways. Furthermore, the degree of DCTPP1 expression was associated with sensitivity to nucleoside DNMTi in correlative and mechanistic studies. Collectively, these results implicate DCTPP1 in cancer cell intrinsic resistance to nucleoside analog DNMTi. The combination of triptolide or its DCTPP1 inhibiting analogs with the existing nucleoside analog DNMTi is therefore ripe for further preclinical and clinical testing in human cancers.

## RESULTS

### A chemical library screen for synergistic combinations with decitabine

We first screened the Johns Hopkins Drug Library, comprised of 3,800 of the estimated 10,000 drugs used in medical practice (Chong et al., 2006; Kamiyama et al., 2013; Platz et al., 2011). We identified those agents that showed >50% growth inhibition (measured by tritiated thymidine incorporation (Platz et al., 2011)) of the prostate cancer cell line DU145 either alone (5 μM of each drug) or in combination with a low dose of decitabine (100 nM) that is well below circulating concentrations of decitabine achieved by clinically used doses (Figure 1A). This narrowed the initial set of compounds to 131 compounds that could have any potential for interaction with decitabine to inhibit DU145 cell growth. After filtering out those compounds that are used only topically, we selected 66 of the remaining compounds to represent a diverse array of compound classes for further analysis in combination studies across a wide dose range of each hit and of decitabine (0 nM to 10 μM each drug). Among these 66 compounds, 24 compounds showed formal synergy with decitabine for DU145 growth inhibition (Chou-Talalay combination index (CI) < 1 at a minimum of 2 decitabine doses and sum of Bliss independence index across the full dose response > 0) (Chou, 2010; Wei et al., 2012) (Figure 1B; Supplementary Figure 1). Among these 24 synergistic hits, there are 6 anti-neoplastic drugs, 6 protein kinase inhibitors, 3 cardiotonic agents, 3 anti-rheumatics and immunosuppressants, 2 antibiotics, 2 mitotic poisons and 2 nucleoside analogs. Interestingly, two of the top 10 synergistic compounds, triptolide and triptonide, are diterpene triepoxides derived from the Thunder God Vine (*Tripterygium wilfordii*) (Figure 1B). Triptolide is reported to be the principal active ingredient from extracts of *T. wilfordii (Titov et al., 2011; Zhou et al., 2012)*, and has been tested in multiple pre-clinical and clinical settings including as an anti-inflammatory, immunosuppressive, anti-fertility, and anticancer agent (Chugh et al., 2012; Zhou et al., 2012). Here, we investigate the role of triptolide in sensitization to nucleoside analog DNMTi.

**Figure 1.**
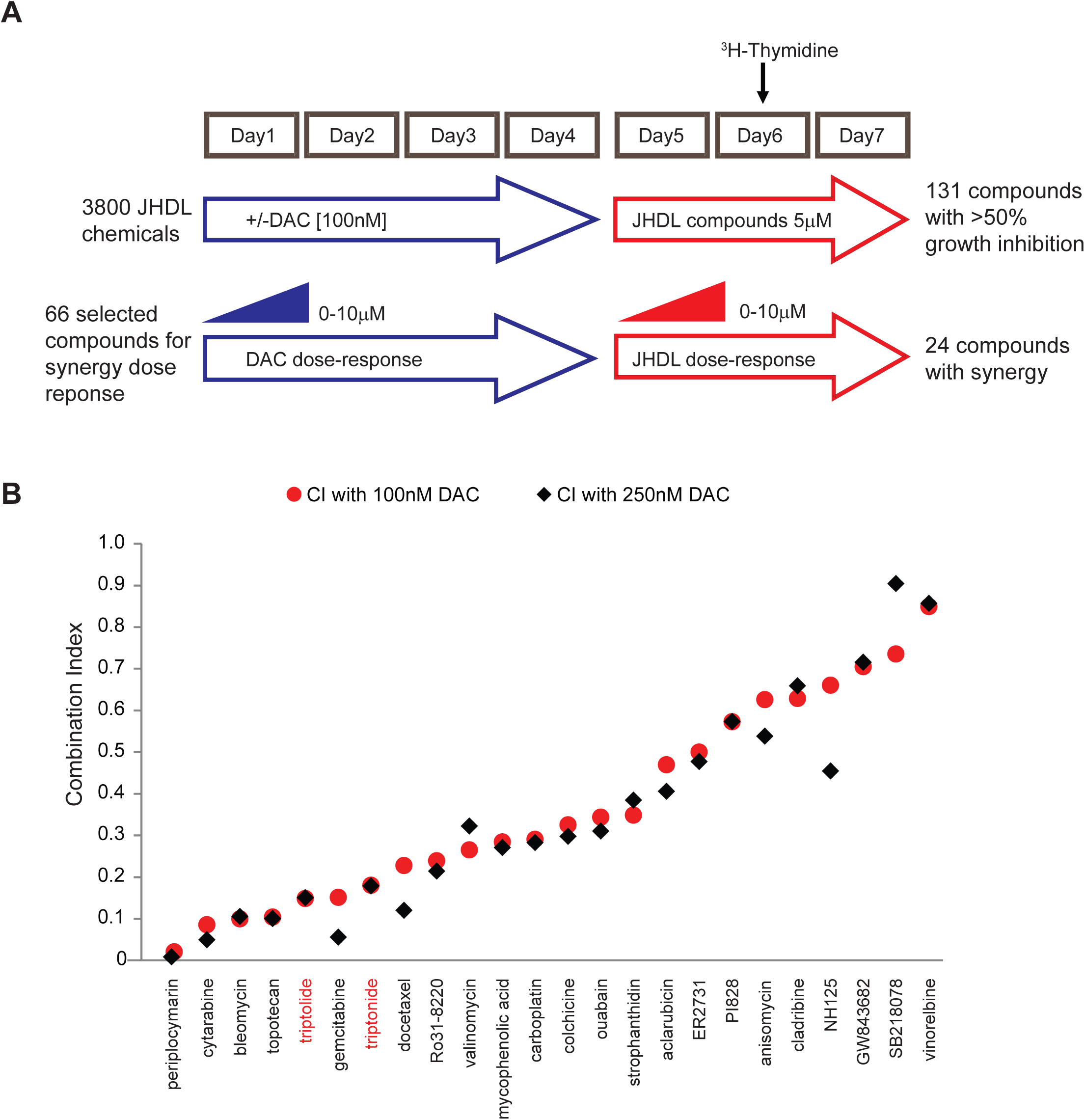
Identification of compounds in the Johns Hopkins Drug Library demonstrating synergy with decitabine for cancer cell growth inhibition. (A). Schematic outline of Johns Hopkins Drug Library (JHDL) screen. DU145 cells were treated with 100nM decitabine or vehicle control for 3 days and then seeded into 96 well-plates for overnight attachment followed by exposure to JHDL compounds (5μM) for 72 hours. The degree of ^3^H-thymidine incorporation (added at day 6 for 18 hours) was measured to evaluate the degree of growth inhibition. 131 compounds showed more than 50% growth inhibition, when given alone or in combination with decitabine, of which 66 compounds were selected for a wide dose-response titration for synergy analysis. Among these, 24 compounds showed synergy with decitabine in inhibiting DU145 cell growth. (B). Scatter plot of 24 synergistic compounds with decitabine, ranked by Combination Index (CI) at a decitabine concentration of 100nM (red circles) or 250 nM (black diamonds). Triptolide and triptonide are shown in red.

### Triptolide synergistically sensitizes cancer cells to nucleoside analog DNMTi *in vitro*

We examined whether the synergy between triptolide and decitabine was more generalizable. When tested alone and in combination using a broad dose response of each agent, triptolide and decitabine exhibited marked synergistic interaction in inhibiting cancer cell growth and/or viability on multiple cancer cell lines including 6 prostate cancer cell lines (DU145, LNCaP, C4-2B, LAPC4, CWR22Rv1, PC3), and 3 leukemia cell lines (K562, CCRF-CEM, MOLT4) (Figure 2A, 2B, Supplementary Figure 2).The combination of triptolide and decitabine also significantly reduced clonogenic survival compared to either drug alone in multiple cancer cell models (Figure 2C). Interestingly, non-malignant RWPE1 prostate epithelial cells and normal hepatocytes in primary culture were largely resistant to the growth inhibitory effects of triptolide and decitabine, both alone and in combination (Figure 2D). This suggests that the combination of triptolide and decitabine may exhibit a favorable therapeutic index.

**Figure 2.**
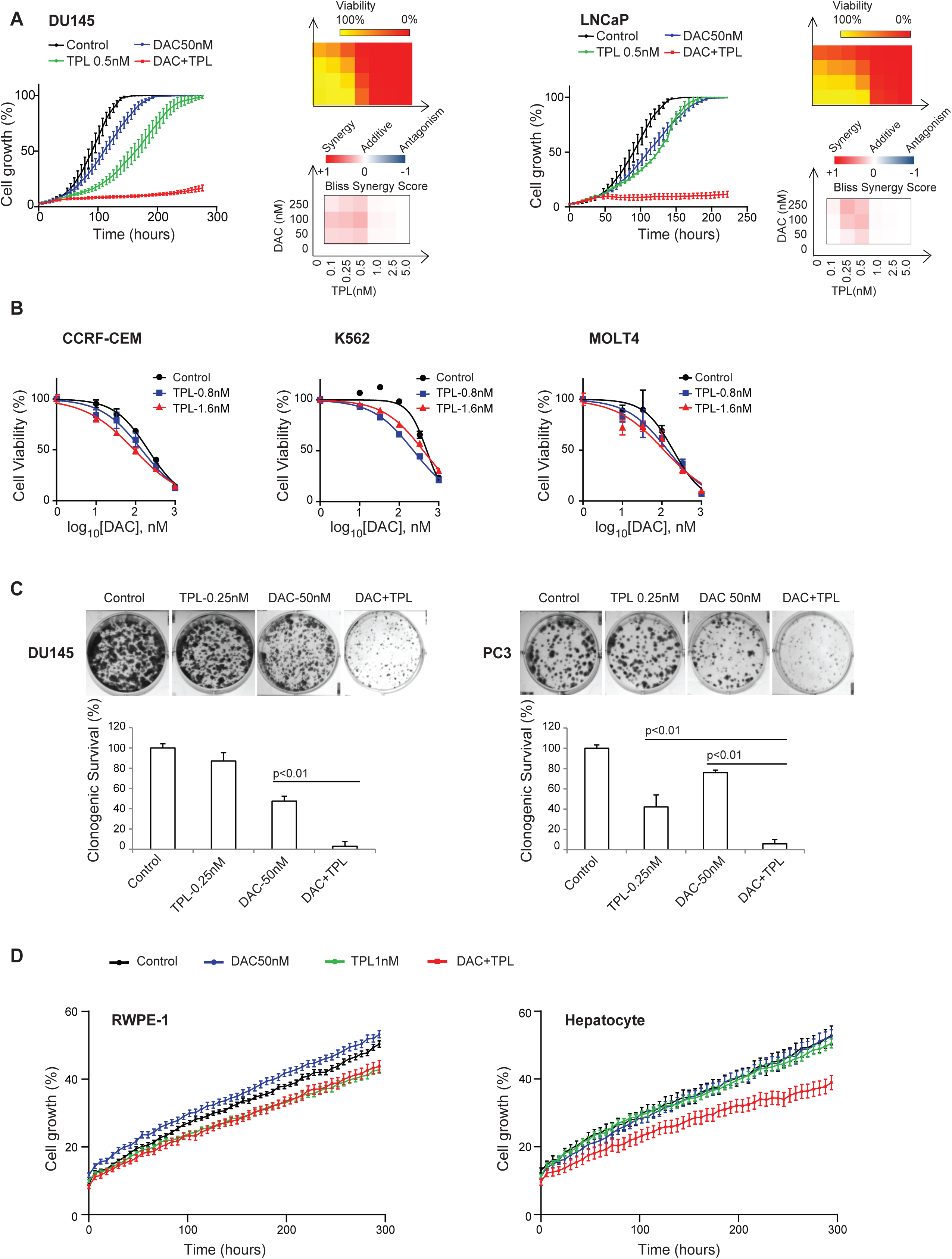
Triptolide and decitabine synergistically and selectively inhibit cancer cell growth, viability, and survival. (A). Inhibition of growth and viability of human prostate cancer cell lines DU145 (left) and LNCaP (right) treated with decitabine and triptolide. Growth curves were measured as the percent of total confluence in each well over time. The mean percent confluence ± SEM for 16 fields in each condition are shown, with measurements taken every 6 hours throughout the course of the growth curve. Viability measurements of cells treated with decitabine and triptolide alone and in combinations across a dose series as indicated are represented in a heatmap scaled as shown in the color legend, with each measurement representing the percent viability with respect to the control treatment. The degree of synergy of combinations of decitabine and triptolide was calculated as the Bliss synergy score across the full dose ranges (values >0 indicate greater than additive effect). For both cell lines, multiple dose combinations exhibited greater than additive effects. (B) Percent viability with respect to the control treatment for human leukemia cell lines treated with decitabine and triptolide in a dose series. Shown are the decitabine alone dose response curve along with the decitabine plus triptolide dose response curves at the two triptolide doses exhibiting the greatest synergy as measured by the Bliss synergy score. Each value represents the mean ± SEM of duplicate treatments.Z (C).Clonogenic survival analysis of human prostate cancer cell lines DU145 (left panel) and PC3 (right panel) treated with vehicle control, decitabine and triptolide alone, and in combination. Quantitation of colonies are normalized to the vehicle treatment and presented as the mean of duplicate wells ± SEM. (D). Growth curve analysis as in (A) for non-malignant RWPE-1 prostate epithelial cells (left) and normal hepatocytes in primary culture (right). At doses that showed synergistic growth inhibition of cancer cell lines (A), decitabine,triptolide and their combination showed minimal growth inhibition of these non-malignant cells.

### Combination of triptolide and decitabine showed enhanced efficacy at well-tolerated doses *in vivo*

To further explore the efficacy and safety of the combination of triptolide and decitabine, we tested each drug alone and in combination in a DU145 prostate cancer xenograft model in athymic nude mice. The combination of decitabine (1 mg/kg doses given i.p. on week 1 on days 1/3/5) and triptolide (0.2 mg/kg doses given each week on days 1-5, 8-12, 15-19) for two 3-week cycles significantly diminished xenograft growth and improved survival compared to triptolide alone, decitabine alone or vehicle control (Figure 3A-C). Interestingly, at these doses in which the combination of triptolide and decitabine showed significantly enhanced efficacy in reducing xenograft growth, we did not observe increased systemic toxicity measured by mouse body weight loss, hematological parameters (WBC, RBC, hemoglobin), liver function (AST, ALT, total protein), and renal function (BUN, creatinine) throughout the treatment course (Figure 3D-G). Taken together, these data suggest that the combination of triptolide and decitabine exhibited favorable efficacy and safety *in vivo*.

**Figure 3.**
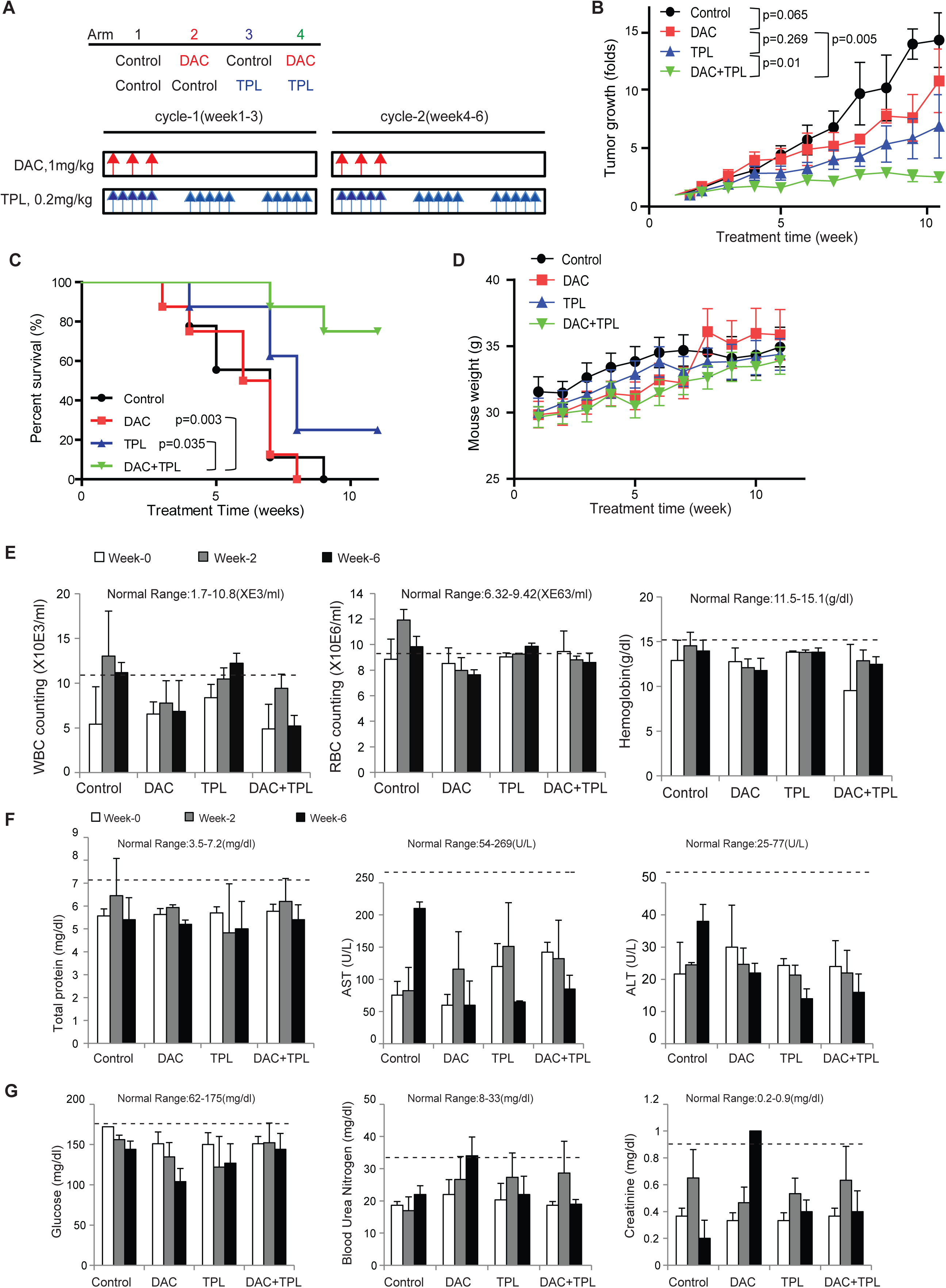
Combination of triptolide and decitabine shows a favorable efficacy and safety *in vivo*. (A). Xenograft study design: NOD/SCID mice bearing DU145 human prostate cancer cell xenografts were divided randomly into 4 groups and then treated with normal saline(Control, Arm1), 1mg/kg decitabine (Arm 2, three 1 mg/kg doses given i.p. on week 1 of two 3-week cycles), 0.2mg/kg Triptolide (Arm3, five 0.2 mg/kg doses given i.p. each week of two 3-week cycles) or in combination (Arm4) upon xenograft tumors reaching an average size of 200 mm^3^. (B). DU145 xenograft tumor growth curve. The tumor volumes were measured every week. The tumor volume in each animal was estimated according to the formula: tumor volume (mm^3^) = 1/2×Length× Width^2^. Combination treatment (Arm4) significantly inhibited the growth of DU145 xenografts compared to single treatment (Arm 2 and Arm 3) and vehicle control (Arm 1). Shown are the mean ± SEM of all xenografted tumors. (C). Kaplan-Meier survival curves of DU145 xenograft mice. Combination of decitabine and triptolide significantly improved progression free survival (time to >5X growth of DU145 xenograft in nude mice) compared to vehicle control (p=0.003), decitabine alone (p=0.003), or triptolide alone (p=0.035). (D). Mouse body weight curve. Mean mouse body weight ± SEM from these experiments are shown. (E, F, G). Changes of hematological (E) parameters (RBC and WBC counting, hemoglobin) and biochemical parameters (Total protein, AST and ALT, glucose, BUN and creatinine) for liver (F) and kidney (G) toxicity monitoring. Each value represents the mean ± SEM of three mice from each treatment arm during treatment (at week 0, 2 and 6). Dashed lines indicate upper limit of the normal range.

### The synergistic interaction of triptolide and decitabine is mediated through inhibition of DCTPP1

We next sought to understand the mechanistic basis for the observed synergy between triptolide and the nucleoside analog DNMTi. To date, a number of direct cellular binding targets for triptolide have been reported, including Xeroderma Pigmentosum B (XPB)/ERCC3 subunit of general transcription factor TFIIH, the dCTP pyrophosphatase DCTPP1, the calcium channel polycystin-2, the membrane protease ADAM10, and the kinase TAK1 partner protein TAB1(Corson et al., 2011; He et al., 2015; Leuenroth et al., 2007; Lu et al., 2014; Soundararajan et al., 2009; Titov et al., 2011). Ensuing mechanistic studies revealed that only XPB among those putative targets is likely the physiological target mediating the antiproliferative activity of triptolide (He et al., 2015; Smurnyy et al., 2014). Among the remaining targets, DCTPP1 appeared to be relevant due to its ability to hydrolyze nucleotide triphosphate in addition to dCTP (Corson et al., 2011; Requena et al., 2014; Song et al., 2015). To understand which of these two direct targets (XPB and DCTPP1) are most responsible for the observed synergy between triptolide and decitabine, we first undertook a chemical biology approach using a series of 6 triptolide analogs that exhibited different potencies for inhibiting DCTPP1 and XPB *in vitro* (Table 1; Supplementary Figure 3A). For each triptolide analog, we determined the IC_50_ for DU145 cancer cell growth inhibition (defined as the dose of the compound needed to reduce cell growth at 150 hours to 50% of vehicle control) with and without combined exposure to decitabine (50 nM). The growth inhibitory potency of each compound alone was significantly correlated to its potency in inhibiting the target XPB (R^2^ =0.882, P=0.006), but not to its potency for DCTPP1 inhibition (R^2^ =0.463, P=0.137; Figure 4A). In a multivariate analysis, the correlation with XPB was preserved (coefficient = 1.11, p = 0.04), while the low trend of correlation with DCTPP1 potency was largely eliminated (coefficient = 0.12, p = 0.76); this suggests that single agent growth inhibition of the triptolide analogs was attributable to XPB targeting, and not DCTPP1 targeting, in agreement with previous findings (He et al., 2015; Smurnyy et al., 2014; Titov et al., 2011).

**Figure 4.**
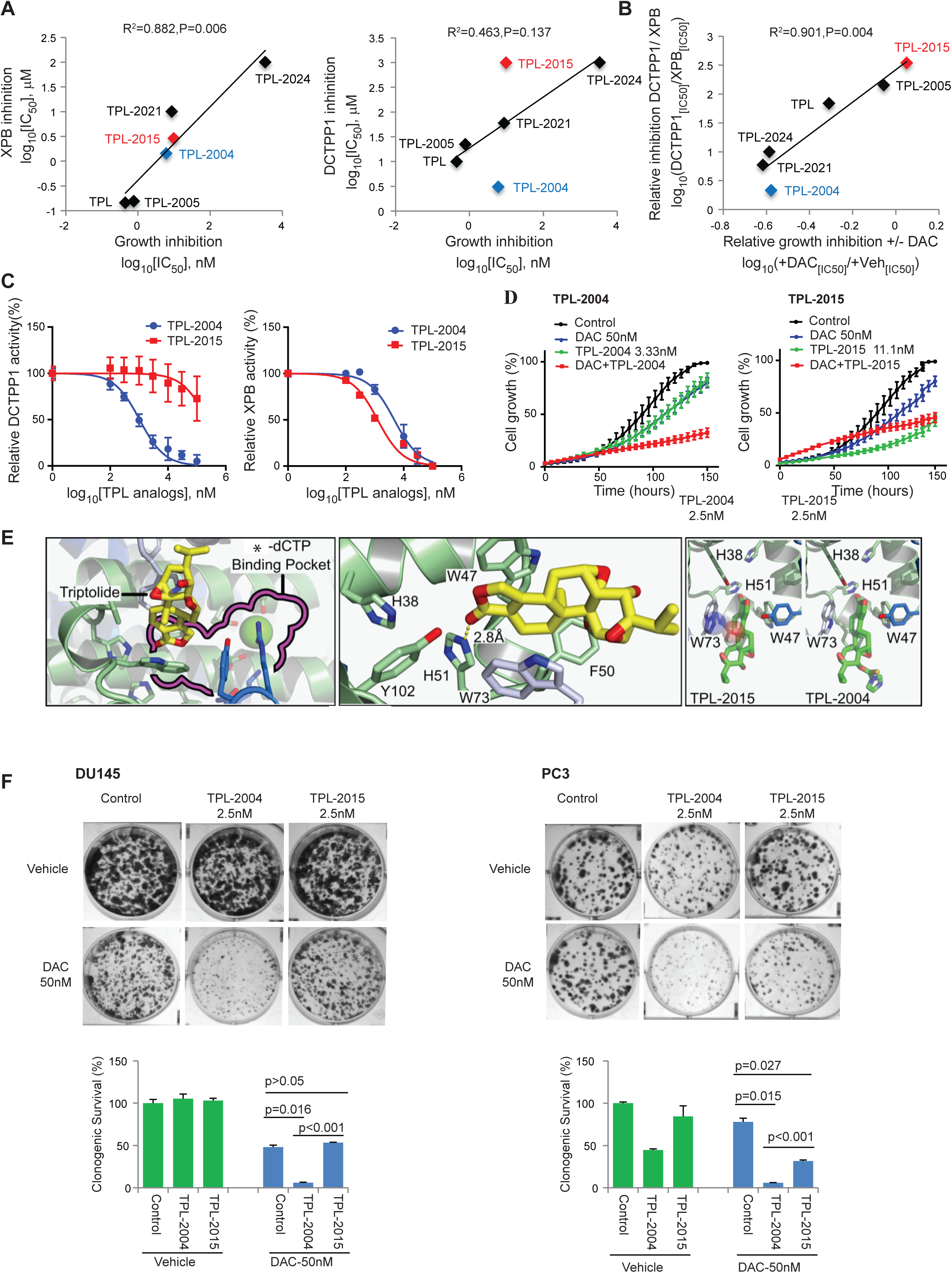
Triptolide analogs with stronger potency for inhibition of DCTPP1 relative to XPB show greater synergy with decitabine. (A). Correlation between potency (log_10_[IC_50_]) of inhibition of DU145 cell growth (measured by growth curve analysis as in Fig 2A) and potency (log_10_[IC_50_]) of inhibition of XPB (left panel) and DCTPP1 (right panel) activity *in vitro* for a series of triptolide analogs. The strong and significant correlation of potency for growth inhibition with inhibition of XPB suggests that triptolide and its analogs mediate single agent growth inhibition via targeting of XPB rather than DCTPP1. (B) Correlation between the log-ratio of [potency of inhibition of DU145 cell growth of each triptolide analog with and without 50 nM decitabine] and the log-ratio of [potency of inhibition of DCTPP1 activity to that of XPB activity]. The strong and significant correlation shows that triptolide analogs with stronger DCTPP1 potency relative to XPB potency exhibit greater enhanced growth inhibition in combination with decitabine. (C). Potency of inhibition of DCTPP1 (left) and XPB (right) *in vitro* activity by the triptolide analogs TPL-2004 and TPL-2015. Both compounds have comparable potency of inhibition of XPB, but TPL-2004 exhibited significantly greater potency of inhibition of DCTPP1 than TPL-2015. Shown are the mean ± SEM of triplicate *in vitro* assays. (D) DU145 growth curve analysis as in Fig 2A with vehicle control, decitabine alone, TPL-2004 or TPL-2015 alone, or each in combination with 50 nM decitabine. Left, TPL-2004; Right, TPL-2015. (E). (Left) The triptolide and nucleotide binding pockets of DCTPP1 overlap. The structure of DCTPP1^(21-130)^ is shown colored in green and blue by monomer subunit. Triptolide is illustrated as yellow sticks, with the dCTP binding pocket as defined from mouse DCTPP1 (PDB#6SQZ) outlined in magenta and a magnesium ion represented as a green sphere. (Center) The triptolide binding pocket is predominantly formed by a hydrophobic sandwich of tryptophan residues W73 and W47, with H51 forming hydrogen bonding contacts with the lactone carbonyl of triptolide. (Right) Triptolide derivatives TPL-2004 and TPL-2015, represented as green sticks, manually overlayed onto the structure of DCTPP1 bound to triptolide. The hydroxyl modification on TPL-2015 (translucent red sphere) would appear to create a steric clash with W73 on DCTPP1 (translucent blue sphere). In contrast, the modifcation on TPL-2004 does not interfere sterically, and appears to close off the triptolide binding pocket. *-dCTP, modified or unmodified dCTP. (F). Clonogenic survival and quantification in DU145 (left panel) and PC3 (right panel) cells treated with vehicle control, decitabine alone, TPL-2004 or TPL-2015 alone, and each in combination with decitabine. Quantitation of colonies are normalized to vehicle control and presented as mean of duplicate wells ± SEM.

**Table 1.**
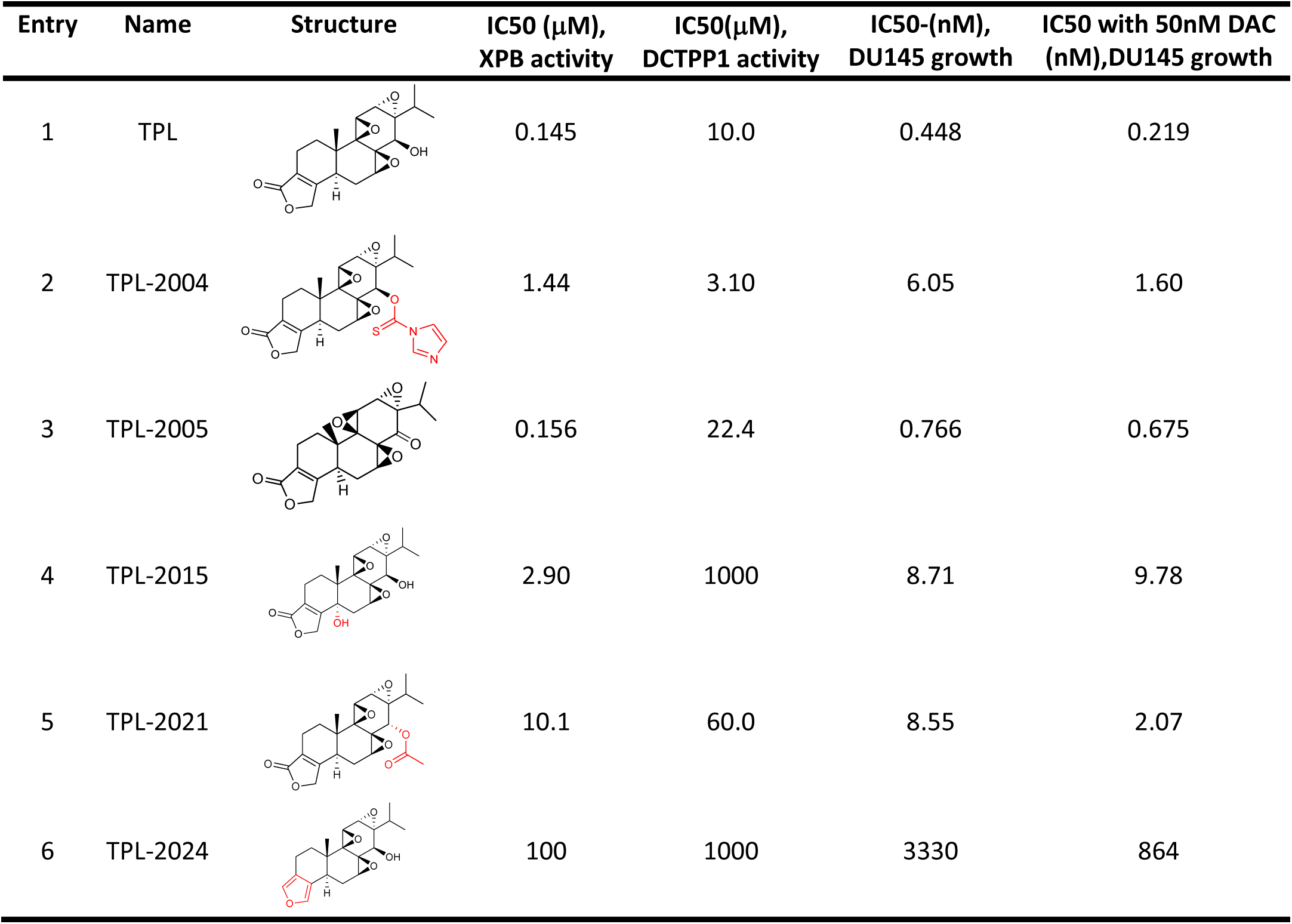
Bioactivities of triptolide (TPL) and its analogs.

| Entry | Name | Structure | IC50 (μM),<br>XPB activity | IC50(μM),<br>DCTPP1 activity | IC50-(nM),<br>DU145 growth | IC50 with 50nM DAC<br>(nM),DU145 growth |
| --- | --- | --- | --- | --- | --- | --- |
| 1 | TPL |  | 0.145 | 10.0 | 0.448 | 0.219 |
| 2 | TPL-2004 |  | 1.44 | 3.10 | 6.05 | 1.60 |
| 3 | TPL-2005 |  | 0.156 | 22.4 | 0.766 | 0.675 |
| 4 | TPL-2015 |  | 2.90 | 1000 | 8.71 | 9.78 |
| 5 | TPL-2021 |  | 10.1 | 60.0 | 8.55 | 2.07 |
| 6 | TPL-2024 |  | 100 | 1000 | 3330 | 864 |

Interestingly, the log ratio of the growth inhibitory IC_50_ for each compound with decitabine to that without decitabine was significantly correlated to the log ratio of IC_50_ for inhibition of DCTPP1 activity vs. XPB activity (R^2^ =0.901, P=0.004; Figure 4B). Thus, the extent to which each compound enhanced growth inhibition with decitabine co-treatment could be attributed to its relative potency for DCTPP1 with respect to XPB. In support of this notion, the degree of synergy between triptolide analogs and decitabine in inhibiting cell growth (measured by the average Bliss synergy score) was correlated with the potency of triptolide analogs in inhibiting DCTPP1 activity more so than that for XPB (Supplementary Figure 3B).

To explore this further, we closely examined the behavior of two triptolide analogs, TPL-2004 and TPL-2015 (Table 1; Figure 4C). Each of these compounds had comparable IC_50_ for inhibition of XPB in activity assays. However, while TPL-2004 had strong potency for inhibition of DCTPP1 activity, comparable to that of the parent compound triptolide, TPL-2015 did not effectively inhibit DCTPP1 (Figure 4C). Likewise, while both TPL-2004 and TPL-2015 showed similar single agent potency for inhibition of DU145 cell growth (6.05 nM and 8.71 nM IC_50_ respectively; Table 1), likely attributable to their comparable potency of inhibiting XPB (Figure 4C), only TPL-2004 and not TPL-2015 showed significantly enhanced growth inhibition with decitabine (Figure 4D). This selective enhancement of growth inhibition by the combination of TPL-2004 with decitabine compared to the combination of TPL-2015 with decitabine was further confirmed by clonogenic survival assays in both DU145 and PC3 cells (Figure 4F).

To define the structural basis for the inhibition of DCTPP1 by triptolide and to understand the differences between triptolide and its analogs, we determined the structure of the catalytic core of human DCTPP1 (residues 21-130) bound to triptolide using X-ray crystallography (Supplementary Table 1). A structure of mouse DCTPP1 has been previously solved in the presence of non-hydrolyzable 5-methyl dCTP (indicated as *-dCTP or Me-dCPNPP in Figure 4E and Supplementary Figure 4 respectively)(Wu et al., 2007), as well as several structures bound to dCTP analogs (Scaletti et al., 2020). The dCTP binding pocket comprised a hydrophobic pyrimidine-binding site and a triphosphate-binding locus lined with charged residues capable of coordinating a catalytic magnesium ion. The pyrimidine-binding pocket is composed of Trp47, Phe50 and Trp73 from an adjacent monomer (human DCTPP1 numbering). The human protein was highly similar structurally to its mouse homolog (Cα rmsd = 0.242Å) (Supplementary Figure 4A), as expected given the sequence similarity between the two proteins (88% identity, 93% similarity for the catalytic core regions). Inspection of the human DCTPP1•triptolide complex revealed that the inhibitor bound in part to the pyrimidine-binding locus (Figure 4E, Supplementary Figure 4C). Trp47 and Trp73 sandwiched the lactone ring and a portion of the diterpine ring of triptolide, while the lactone carbonyl of the compound formed hydrogen bonds to His51, a residue also seen to engage the 4-amino group of dCTP. Phe50 also made close contact to the diterpene ring of triptolide, while residues Tyr102 and Tyr129 formed an additional adjacent monomer to further define the hydrophobic binding pocket.

Triptolide contains three epoxides. The inhibition of XPB by triptolide has been attributed to covalent adduct formation between a cysteine in the protein and one of the three epoxides (Titov et al., 2011). In the DCTPP1•triptolide complex, the binding pocket contained no cysteines or serines and no covalent adducts were observed (Figure 4E, Supplementary Figure 4B). The structure of the complex also explained the disparate activities of analogs TPL-2004 and TPL-2015. The Hydroxyl substitution of TPL-2015 likely disrupted the hydrophobic sandwich formed by W47 and W73, accounting for the >100-fold difference in IC_50_ of this derivative for DCTPP1 compared to triptolide (Figure 4E). By comparison, TPL-2004 was functionalized with an imidazole group linked through a thio-enol ester to a hydroxyl group on triptolide distal to its lactone ring. This imidazole substituent likely closed off the hydrophobic binding pocket exploited by triptolide (Figure 4E), accounting for its observed ∼3-fold improvement in IC_50_ (Table 1).

To further determine the mechanism of synergy between decitabine and triptolide, we carried out additional genetic experiments to complement the chemical biology approaches described above. CRISPR/CAS9 mediated genetic disruption of DCTPP1 led to sensitization of DU145 and PC3 cells to single agent decitabine (Figure 5A), and greatly abolished the synergy between decitabine and triptolide (P<0.001, Fig 5B, 5D). However knocking out XPB in DU145 and PC3 did not significantly impact the sensitivity to decitabine or the synergy between decitabine and triptolide (Supplementary Figure 5A-C). On the other hand, forced expression of DCTPP1 in DU145 and PC3 cells led to significant desensitization to single agent decitabine (Figure 5A), and enhanced the synergy between decitabine and triptolide in inhibiting cancer cell proliferation across a dose response matrix (Figure 5C, 5E). In contrast, forced expression of XPB in DU145 and PC3 cells had little impact on sensitivity to single agent decitabine (Supplementary Figure 5D).

**Figure 5.**
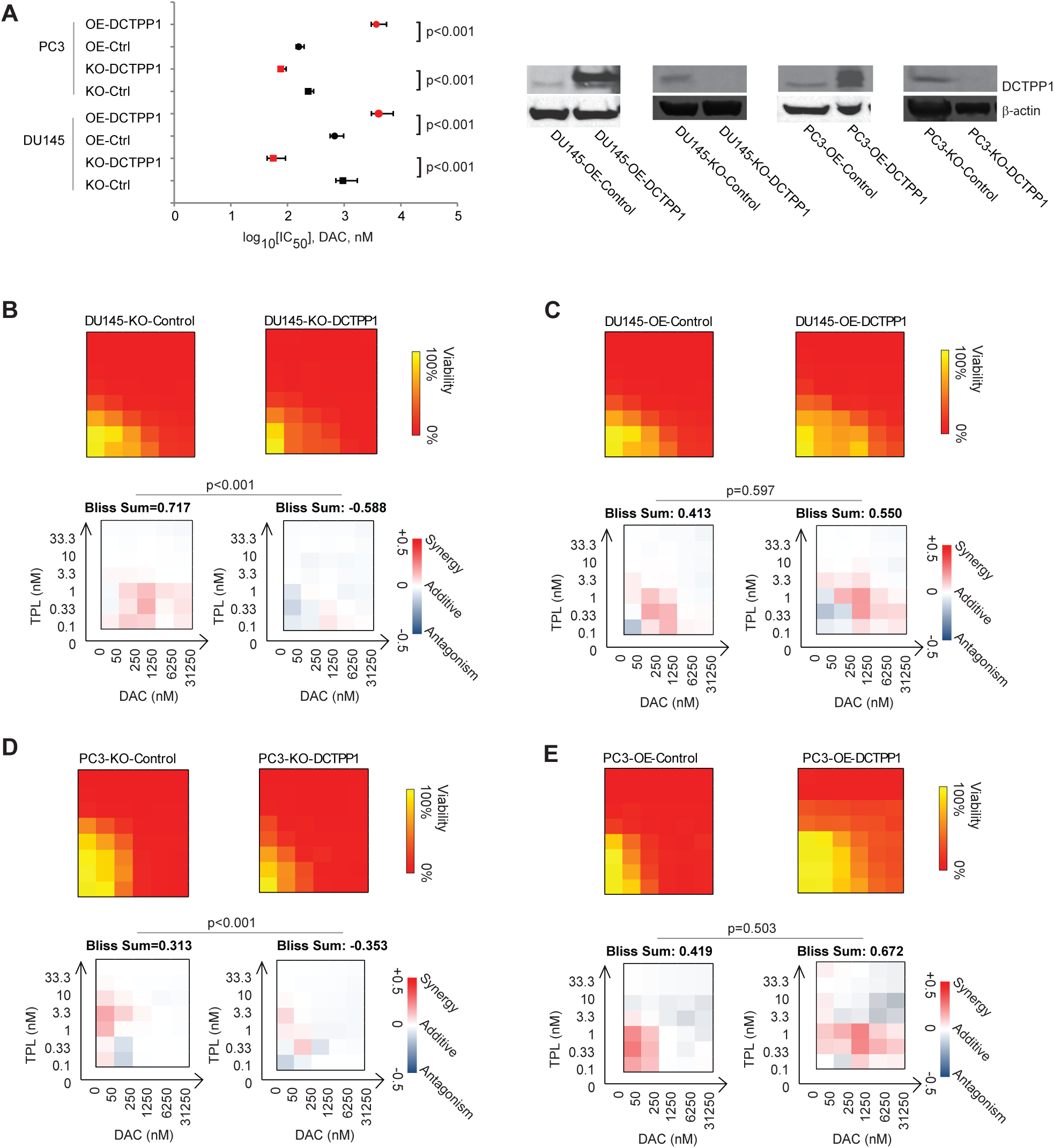
The synergy of triptolide and decitabine is dependent on DCTPP1. (A). Cellular sensitivity to decitabine is dependent on DCTPP1 levels. Human prostate cancer cell lines PC3 and DU145 were transduced with CRISPR-Cas9 system with DCTPP1 guide-RNA to knockout DCTPP1 (KO-DCTPP1), or non-targeting guide-RNA (KO-Control), or DCTPP1 overexpression lentivirus (OE-DCTPP1), or empty lentivirus (OE-Control). Knockout and overexpression of DCTPP1 was confirmed by immunoblotting in right panels. Potency of inhibition of growth and viability of the transduced isogenic cell lines was measured across a dose range of decitabine. Overexpressing DCTPP1 significantly desensitized the cells to decitabine (significant right shift of log_10_[IC_50_]); while knocking out DCTPP1 significantly sensitized the cells to decitabine (significant left shift of log_10_[IC_50_]). (B,C,D,E). The synergy between decitabine and triptolide is dependent on DCTPP1 expression levels in isogenic DU145 and PC3 cells. Viability measurements of DU145 and PC3 cells treated with decitabine and triptolide alone and in combinations across the indicated dose series are represented in heatmaps scaled as shown in the color legend, with each measurement representing the percent viability with respect to the control treatment. The degree of synergy of combinations of decitabine and triptolide was calculated as the Bliss synergy score across the indicated dose range. Knocking out DCTPP1 in DU145 (B) and PC3 (D) cells showed no overall synergy across serial dose combinations of decitabine and triptolide (Bliss values <0 indicate less than additive effect). Both DU145 (C) and PC3 (E) cells with DCTPP1 overexpression retained slightly better overall synergy across serial dose combinations of decitabine and triptolide (Bliss values >0 indicate greater than additive effect).

Taken together, these findings suggest that inhibition of cell growth by single agent triptolide and its analogs is likely due to their ability to inhibit the target XPB, while the observed synergy between triptolide and its analogs with decitabine for inhibition of cancer cell growth and survival is attributable to inhibition of DCTPP1.

### Defining the mechanism of synergy of DNMT and DCTPP1 inhibition

Since DCTPP1 was the major target for triptolide and its analogs for synergy with decitabine in inhibiting cancer cell growth and survival, we next explored the underlying mechanisms. DCTPP1 encodes the deoxycytidine triphosphate pyrophosphatase, which is involved in balancing levels of deoxycytidine triphosphate (dCTP) and deoxycytidine monophosphate (dCMP) in cells. DCTPP1 has also been shown to hydrolyze pyrophosphate from non-natural and modified dCTP, including 5-halogenated, 5-formyl, 5-methyl, 5-hydroxymethyl, more so than dCTP itself, potentially acting to more selectively prevent the incorporation of these modified dCTP nucleotides into genomic DNA (Corson et al., 2011; Requena et al., 2014; Song et al., 2015).

The convergent mechanism of action of the nucleoside analog DNMTi, including decitabine, azacitidine, and guadecitabine, involves cellular conversion to 5-aza-2’-deoxycytidine triphosphate (5-aza-dCTP) and incorporation into the genome during DNA synthesis, triggering trapping and degradation of the DNA methyltransferase enzymes and passive loss of methylation across DNA replication (Ghoshal et al., 2005; Patel et al., 2010; Yang et al., 2010). We hypothesized that DCTPP1 may have pyrophosphatase activity on 5-aza-dCTP, thereby preventing its incorporation into genomic DNA, presenting a cell-intrinsic resistance mechanism to nucleoside analog DNMTi.

To begin testing this hypothesis, we examined the enzymatic activity of DCTPP1 for hydrolysis of pyrophosphate from 5-aza-dCTP and dCTP. These analyses revealed that DCTPP1 had slightly increased catalytic efficiency (Kcat/Km) for 5-aza-dCTP compared to dCTP (Supplementary Figure 6A). We also observed that triptolide can inhibit DCTPP1 pyrophosphatase activity for 5-aza-dCTP with an IC_50_ similar to that for dCTP (14.7 μM and 8.49 μM respectively) (Supplementary Figure 6B). Additionally, we showed that decitabine did not inhibit DCTPP1 pyrophosphatase activity (Supplementary Figure 6C). These data suggest that DCTPP1 may reduce 5-aza-dCTP availability for incorporation into genomic DNA, and that triptolide can increase the availability of 5-aza-dCTP incorporation by inhibiting DCTPP1 pyrophosphatase activity.

Consistent with this notion, co-treatment with decitabine and triptolide led to enhanced 5-aza-dC incorporation into genomic DNA compared to treatment with decitabine alone as measured by a liquid chromatography/tandem mass spectrometry assay (Figure 6A, 6B)(Anders et al., 2016). Similarly, compared to that in control DU145 cells, the level of 5-aza-dC incorporation into genomic DNA after decitabine treatment was significantly reduced in DCTPP1-overexpressing cells (Supplementary Figure 7A), and was significantly increased in DCTPP1-knockout cells (Supplementary Figure 7B). Intriguingly, this enhanced genomic incorporation of 5-aza-dC by combination decitabine and triptolide treatment translated to significantly enhanced trapping and degradation of all three DNA methyltransferases (DNMT1, DNMT3A, and DNMT3B) in the cells (Figure 6C, Supplementary Figure 8A-B) as well as a greater degree of global DNA demethylation (Figure 6D,E), compared to decitabine alone, which could only lead to reduction of DNMT1 levels. This enhanced demethylation was also observed when decitabine was combined with the DCTPP1-inhibiting triptolide analog TPL-2004, but not with TPL-2015 (Figure 6F).

**Figure 6.**
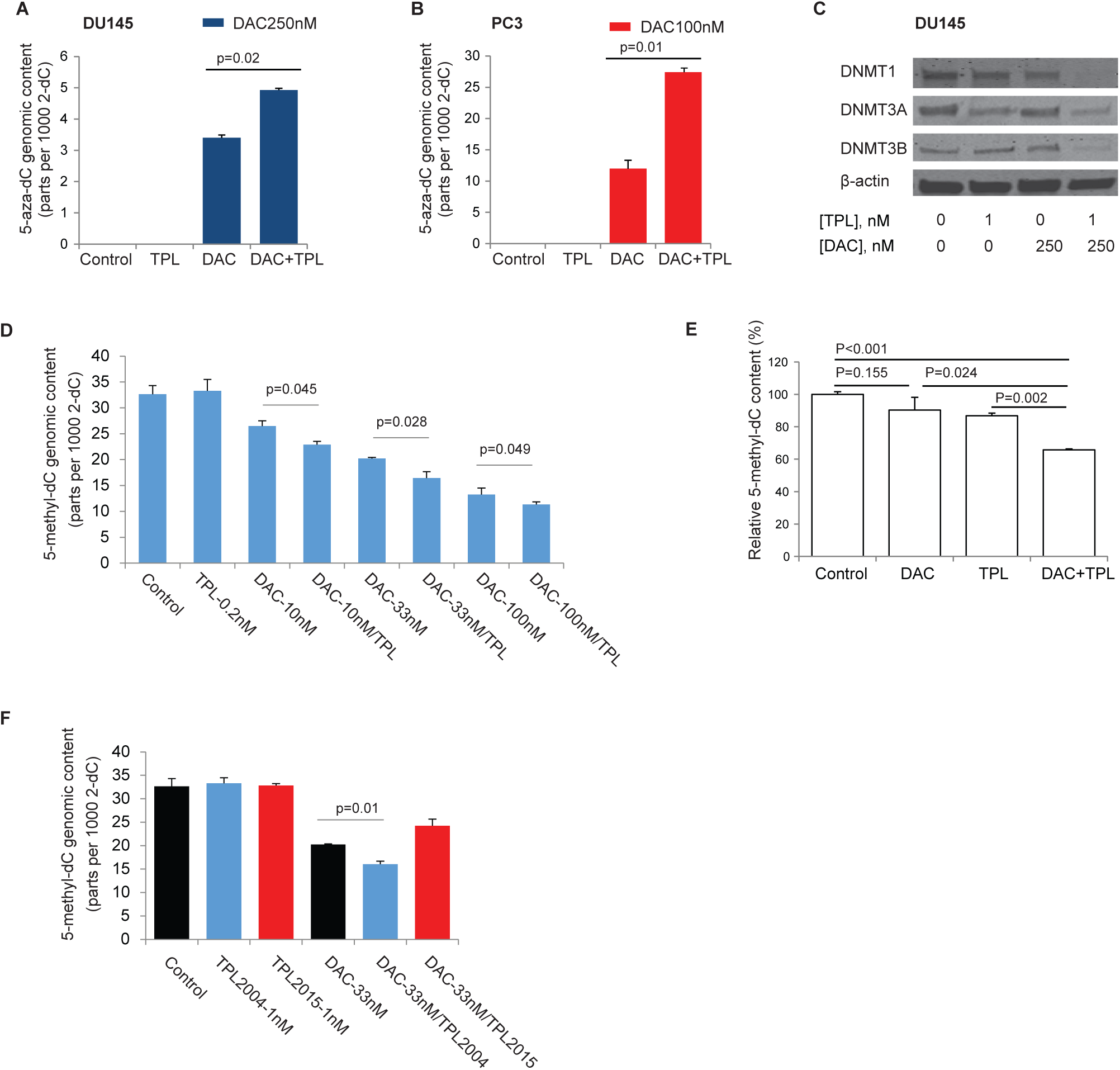
DCTPP1 inhibition by triptolide enhances the action of nucleoside analog DNMT inhibitors. (A, B). 5-aza-dC incorporation into genomic DNA, determined by LC-MS/MS, in DU-145 (A) and PC3 (B) cells treated with vehicle control, triptolide or decitabine alone or in combination. The mean of the amount of 5-aza-dC per 1000 2-dC in genomic DNA ± SEM of duplicate treatments is shown. Triptolide treatment significantly enhanced the decitabine incorporation into genomic DNA in DU145 and PC3 cells. (C). The combination treatment of triptolide and decitabine enhanced DNMT loss. Western Blot of DNMT1, DNMT3A, DNMT3B in DU145 cells treated with vehicle control, decitabine (250 nM) or triptolide (1 nM) alone or in combination for 24 hours. β-actin blotting is shown as protein loading control. (D) Genomic 5-methyl-deoxycytidine content by LC-MS/MS in DU145 cells treated *in vitro* with vehicle control, decitabine or triptolide or both at indicated doses. Each value represents the mean ± SEM of 5-methyl-deoxycytidine content from triplicate measurements at 5 days from the start of treatment. The combination treatment of triptolide and all tested doses of decitabine significantly lowered the levels of 5-methyl-deoxycytidine content in genomic DNA compared to each dose of decitabine alone. (E) Genomic 5-methyl-deoxycytidine content by LC-MS/MS in DU145 xenograft tumors treated *in vivo* with vehicle control, decitabine or triptolide or both. Each value represents the mean ± SEM of 5-methyl-deoxycytidine content from two xenografted tumors as a percent of that measured in xenografted tumors from vehicle control treated animals. (F) Genomic 5-methyl-deoxycytidine content by LC-MS/MS in DU145 cells treated *in vitro* with vehicle control, triptolide analogs (TPL-2004 or TPL-2015), decitabine or triptolide or both. Each value represents the mean ± SEM of 5-methyl-deoxycytidine content from triplicate measurements at 5 days from treatment starting.

### Combination of DNMTi with triptolide enhances the epigenetic effects of decitabine in surviving cells

Prior work has suggested that high doses of nucleoside DNMTi lead to significant cytotoxicity without mediating significant epigenetic effects in the surviving cells; in contrast, low doses of nucleoside DNMTi mediated significant DNA demethylation and long term epigenetic reprogramming of gene expression but without significant cytotoxicity (Blagitko-Dorfs et al., 2013; Chiappinelli et al., 2015; Leonard et al., 2014; Scandura et al., 2011; Tsai et al., 2012; Ye et al., 2016). We hypothesized that the combination of triptolide (or its DCTPP1-inhibiting analogs) with decitabine could allow significant cancer cell cytotoxicity, while also mediating DNA demethylation and epigenetic reprogramming in any surviving cells. To test this hypothesis, we exposed DU145 cells to vehicle controls, triptolide (0.5 nM) or decitabine (33 nM) alone at low doses, or the combination, for 3 days, replaced with drug free media, and assessed the surviving cells for genome-wide DNA methylation (using the Infinium EPIC microarray platform) and gene expression (using RNA-seq) patterns over a time course at 5 days and 20 days post initial exposure. As expected, the combination of triptolide and decitabine showed significant inhibition of growth/survival at 5 days while each drug alone had minimal effect. Also as expected, low dose decitabine alone, but not triptolide alone, led to marked genome-wide DNA demethylation (Supplementary Figure 9A). Interestingly, the combination of low dose triptolide and decitabine also showed marked genome-wide DNA demethylation, and enhanced the degree of DNA demethylation compared to low-dose decitabine alone to some extent. Moreover, this pattern of genome-wide DNA demethylation was retained in the surviving cells even at 20 days post-exposure (Supplementary Figure 9B). Consistent with these in vitro findings, in the long-term *in vivo* experiment, mice treated with the combination of decitabine and triptolide showed significantly greater loss of 5-methyl-deoxycytidine (5mdC) in the residual tumor genomic DNA compared to that in mice treated with decitabine alone (Figure 6E).

Remarkably, in terms of gene expression, we also observed that the genes significantly induced by decitabine alone (Benjamini-Hochberg adjusted p (padj) < 0.001, log_2_(fold change) > 0.5), across the board, were subtly but consistently further upregulated by the combination of decitabine and triptolide (p <0.001; Figure 7A, Supplementary Figure 10). As an example, the long noncoding RNA H19, which is known to exhibit loss of imprinting and DNA methylation mediated silencing in DU145 cells, was highly induced by decitabine alone, and was further upregulated by the combination of triptolide and decitabine, which was confirmed by qRT-PCR (Figure 7B). Deep bisulfite sequencing of the previously characterized imprint control region of H19 (Barletta et al., 1997; Ito et al., 2013) showed that the combination of decitabine and triptolide led to a greater fraction of completely demethylated alleles compared to decitabine alone (Figure 7C). Consistent with prior reports (Chiappinelli et al., 2015; Roulois et al., 2015), in our experimental system, decitabine strongly induced expression of endogenous retroviruses, cancer testis antigens, and interferon response genes (Figure 7D-F). The combination of triptolide and decitabine further enhanced induction of these genes compared to decitabine alone (Figure 7D-F).

**Figure 7.**
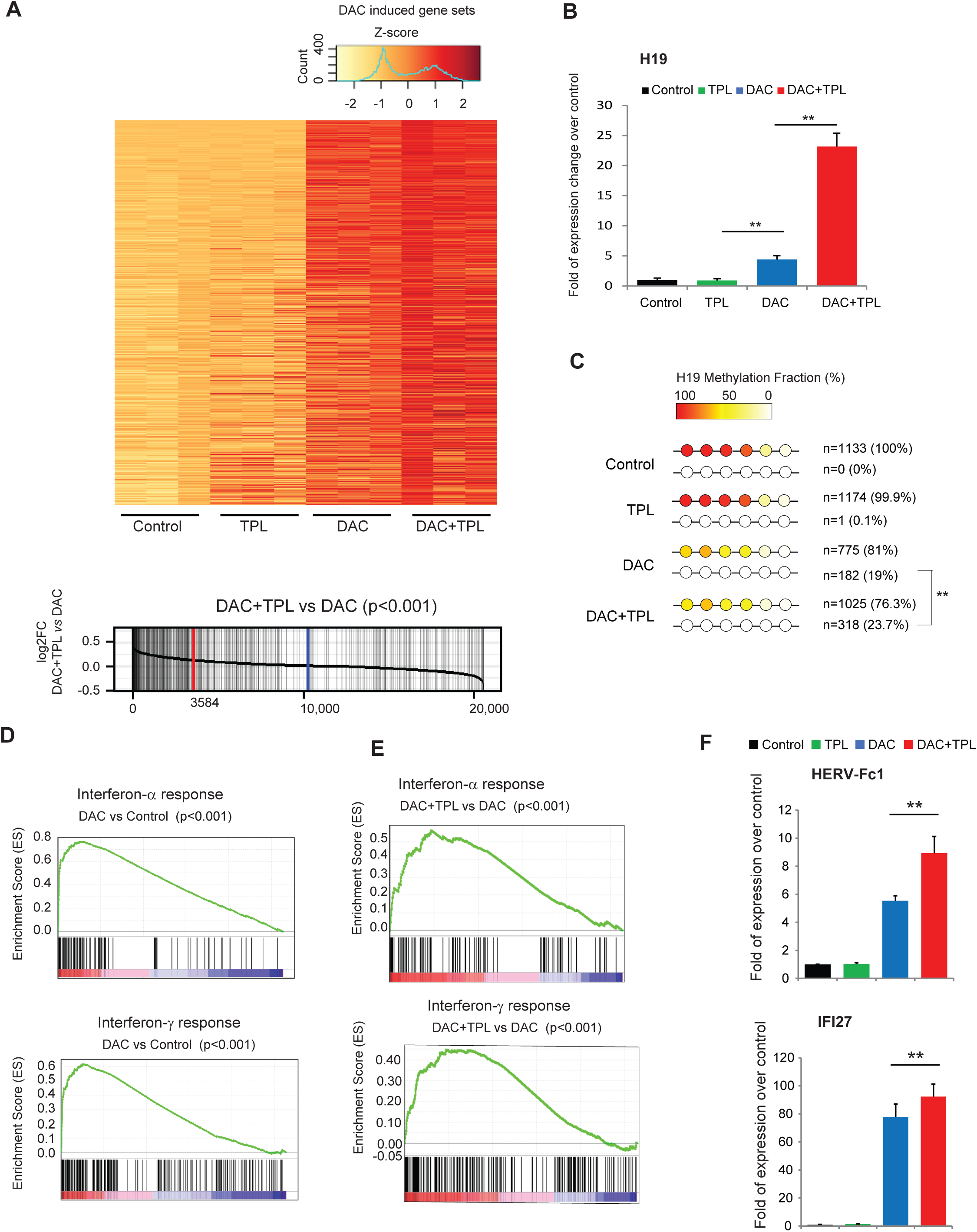
The combination treatment of triptolide and decitabine enhances the epigenetic effects of decitabine. (A). Decitabine induced gene sets (n = 704, DAC/DAC+TPL vs Control,log2 Fold of Change >0, P < 0.001) are further upregulated by treatment with the combination of triptolide and decitabine. (Upper panel) Heatmap representing day-20 gene expression profiles of triplicate samples of DU145 cells treated with vehicle control (Control), 2nM triptolide alone (TPL), 33nM decitabine alone (DAC), or in combination (DAC+TPL). (Lower panel) Distribution of decitabine induced genes across the whole transcriptome in combination treatment of triptolide and decitabine. Each vertical line represents a decitabine upregulated gene (n=704, DAC+TPL vs DAC, p<0.001, Wilcoxon rank sum test), ranked by log_2_ fold of change of combination treatment of decitabine and triptolide over decitabine alone (horizontal line, P < 0.001, Wilcoxon rank sum test). Red vertical line represents the median for decitabine induced genes; Blue vertical line represents median of whole transcriptome. 525 of 704 decitabine induced genes were further upregulated by triptolide and decitabine combination treatment. (B). The long noncoding RNA H19, is one of the top decitabine induced genes and also upregulated in decitabine and triptolide combination treatment. qRT-PCR validated there were significantly higher mRNA expression of H19 compare to decitabine alone, triptolide alone or vehicle control.(**: p<.001). (C). Bisulfite sequencing analysis of H19 imprint control region in the representative samples at day-20 after the indicated treatment. The combination of decitabine and triptolide led to a greater fraction of completely demethylated alleles compared to decitabine alone (**: p<.001, Binomial test). Data shown are the numbers and patterns of methylated and unmethylated clones and their percentages. Color scale shows methylation fraction at each CpG sites. (D,E). DU145 cells treated with decitabine alone (D) are enriched for increased expression of interferon-α response genes (upper panel, p < 0.005, GSEA) and interferon-γ response genes (lower panel, p < 0.05, GSEA). The combination treatment of triptolide and decitabine (E) further enhanced induction of these genes compared to decitabine alone. (F). qRT-PCR validation of endogenous retrovirus gene HERV-FC1 (upper panel) and interferon response genes IFI27 (lower panel). Data shown are represented as mean ± SEM of three biological replicates at day-15 after indicated treatment, y axis is represented as fold change over vehicle control treatment. **: p<0.001.

Thus, compared to decitabine alone, the combination of triptolide and low dose decitabine captures the most desired effects of both low-dose and high-dose nucleoside DNMTi, in that it leads to both selective cancer cell cytotoxicity while also mediating, and even enhancing, the long-lived epigenetic effects of DNA demethylation and gene re-expression.

### DCTPP1 inhibition by triptolide overcomes a cell intrinsic resistance mechanism to nucleoside analog DNMTi

These data implicate DCTPP1 as a cell-intrinsic factor in mediating primary resistance to nucleoside analog DNMTi by preventing their incorporation into genomic DNA. To further investigate this notion, we assessed whether the degree of incorporation of decitabine into genomic DNA in a panel of 16 human cancer cell lines (5 lung cancer, 4 breast cancer, 4 melanoma, 2 prostate cancer and 1 ovarian cancer) was associated with the level of expression of DCTPP1 or with any other genes from the “pyrimidine metabolism” pathway curated in the Kyoto Encyclopedia of Genes and Genomes (KEGG; Figure 8A)(Yang et al., 2010). Interestingly, there was a significant inverse correlation with 5-aza-dC incorporation and DCTPP1 expression levels (R^2^ =0.493, p=0.003, Figure 8A-C) (Cerami et al., 2012; Gao et al., 2013), and this correlation was preserved even after normalizing to differences in the rate of proliferation between cell lines as measured by tritiated thymidine incorporation (Supplementary Figure11A). This correlation with DCTPP1 expression was the top-ranked correlation among all genes in the “pyrimidine metabolism” KEGG pathway, including those that are known to be directly involved in the cellular uptake and metabolism of decitabine (Figure 8A,B). Conversely, there was no significant correlation between 5-aza-dC incorporation and the level of expression of XPB, the other known major direct target of triptolide (Supplementary Figure 11B). Furthermore, the growth inhibitory potencies of decitabine were also significantly correlated with the DCTPP1 expression levels among a panel of 18 cancer cell lines (R^2^ =0.238, p=0.0399, Figure 8D). In cell lines with intermediate to high DCTPP1 expression (DU145, PC3, A549 and MCF7), CRISPR/CAS mediated disruption of DCTPP1 significantly sensitized (reduced IC_50_) the cells to single agent decitabine compared to those of the isogenic control cells (Figure 5A, 8E, Supplementary Figure 12A, 12C). On the other hand, in cell lines having low to intermediate DCTPP1 expression (MDA-MB-231, DU145 and PC3), overexpressing DCTPP1 conferred resistance (increased IC_50_) to single agent decitabine compared isogenic control cells (Figure 5A, 8F, Supplementary Figure 12A,12B).

**Figure 8.**
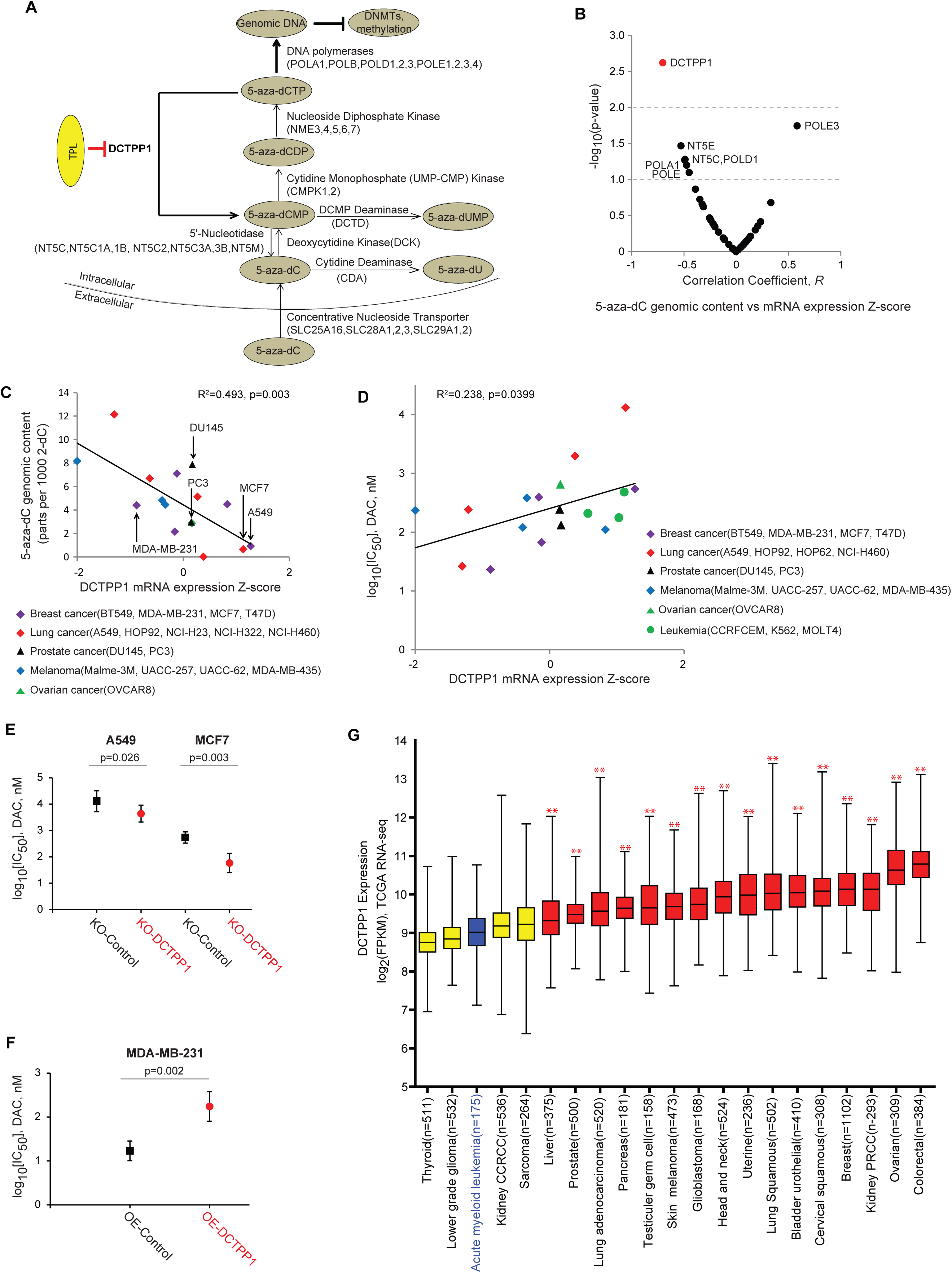
DCTPP1 expression leads to reduced 5-aza-dC incorporation and decitabine potency. (A). A schematic representation of the enzymes (with HUGO gene symbols of encoding genes in parentheses) curated in the “pyrimidine metabolism” KEGG pathway that are directly involved in decitabine uptake, metabolism and incorporation. Our proposed model of synergy between decitabine and triptolide is shown with bolded arrows: by inhibiting DCTPP1 (shown in bolded red arrow), triptolide prevents pyrophosphate cleavage of 5-aza-deoxycytidine triphosphate (5-aza-dCTP), resulting in greater incorporation of 5-aza-dC into genomic DNA. This would trigger increased DNMT trapping/degradation, greater hypomethylation, and synergistic cytotoxicity in vitro and in vivo (shown in bolded black arrow). 5-aza-dC, 5-aza-2’-deoxycytidine; 5-aza-dU, 5-aza-2’-deoxyuridine; 5-aza-dCMP, 5-aza-2’-deoxycytidine monophosphate; 5-aza-dUMP, 5-aza-2’deoxyuridine monophosphate; 5-aza-dCDP, 5-aza-2’deoxycytodine diphosphate; 5-aza-dCTP, 5-aza-2’-deoxycytidine triphosphate; DNMTs, DNA methyltransferases. (B). A plot of the correlation coefficient versus the [–log_10_(p-value)] for the correlation between 5-aza-dC incorporation into genomic DNA and the level of expression for each gene from the Kyoto Encyclopedia of Genes and Genomes (KEGG) “pyrimidine metabolism” pathway that would be directly involved in decitabine uptake, metabolism/activation, and DNA incorporation. The Human Genome Organization (HUGO) gene symbols for the top-ranked genes (with p < 0.1) are indicated. Among these genes, only DCTPP1 showed a significant (p<0.01) correlation with the level of incorporation of 5-aza-dC into genomic DNA. (C). The level of incorporation of 5-aza-dC into genomic DNA after two days of exposure to 250 nM decitabine is significantly inversely correlated with the level of expression of DCTPP1 in the panel of 16 cancer cell lines. The cells selected for genetic manipulation are indicated with arrows. (D). Decitabine potency (log_10_(IC50)) is significantly correlated with the level of DCTPP1 expression in a panel of 18 cancer cell lines (4 lung cancer, 4 breast cancer, 4 melanoma, 3 leukemia, 2 prostate cancer and 1 ovarian cancer). (E,F). Genetic disruption of DCTPP1 significantly sensitized A549 and MCF7cells to decitabine (E), while overexpressing DCTPP1 in MDA-MB-231 cells conferred significant resistance to decitabine (F). Data represent log_10_ [IC_50_, DAC (nM)] and 95% confidence intervals. (G). DCTPP1 expression levels in acute myeloid leukemia are significantly lower than those in most solid tumors. The DCTPP1 mRNA expression levels among the major cancer types in The Cancer Genome Atlas (TCGA, http://www.cbioportal.org/) were plotted and each cancer type was ranked by its median DCTPP1 expression level. The box of the box-and-whisker plots showed the median and interquartile range, and the whiskers represent the minimum and maximum log_2_(RPKM) value (reads per kilobase per million). DCTPP1 mRNA expression levels in acute myeloid leukemia (blue) were significantly lower than those in most solid tumor samples(red), including liver, lung, testis, prostate, brain glioblastoma, head and neck, bladder, breast, ovarian, cervical and colorectal cancers (**, p<0.001). There is no significant difference in DCTPP1 mRNA expression between AML and the remaining four cancer types (yellow). CCRCC, clear cell renal cell carcinoma; PRCC, papillary renal cell carcinoma.

Nucleoside analog DNMTi, including decitabine, azacitidine, and guadecitabine, have shown clinical efficacy in myelodysplastic syndromes (MDS) and acute myeloid leukemia (AML) (DiNardo et al., 2020; Issa et al., 2015; Jones and Baylin, 2002; Kwag et al., 2022; Yang et al., 2010), but have been disappointing as single agents in many solid organ cancers. Interestingly, an analysis of DCTPP1 mRNA expression levels in multiple cancer types from the TCGA showed that the level of DCTPP1 expression in AML (n = 175) is significantly lower than those in the majority of major solid organ cancers (Figure 8G). Furthermore normal bone marrow and liver tissues had relatively low DCTPP1 expression (Supplementary Figure 13), suggesting that there may not be an enhanced hematopoietic or liver toxicity of combination DNMT and DCTPP1 inhibition compared to DNMT inhibition alone. We can speculate that the relatively low level of DCTPP1 expression in bone marrow and hematological malignancies compared to most normal tissues and solid tumors respectively may be partially responsible for the observed hematological side effects and clinical sensitivity to nucleoside analog DNMTi in hematological malignancies like MDS and AML.

We further tested whether inhibition of DCTPP1 by triptolide may have more broad implications for sensitization to several classes of cytidine nucleoside analog drugs whose mechanism of action involves incorporation into DNA. The other FDA approved nucleoside DNMTi azacitidine can be metabolized and incorporated into both RNA (∼80-90%) and DNA (∼10-20%); however, the major mechanism of action in MDS/AML and other cancers is thought to arise from the conversion (through ribonucleotide reductase and other enzymes) to 5-aza-dCTP and incorporation into DNA, thereby exerting both epigenetic demethylation and DNA damage responses (Baylin, 2005; Kiziltepe et al., 2007; Palii et al., 2008). We found that, similar to decitabine, co-treatment of DU145 cells with azacitidine and triptolide or its DCTPP1-inhibiting analog TPL-2004 showed synergistic cytotoxicity, enhanced 5-aza-dC incorporation into genomic DNA and global demethylation compared to treatment with azacitidine alone (Supplementary Figure 14A-B, Supplementary Figure 15A-B). Since DCTPP1 does not have enzymatic activity on ribonucleosides, these data provide further confirmation that the major mechanism of action of azacitidine is through incorporation in DNA and not RNA. In further support of this notion, we found that across a panel of 15 cell lines, the degree of 5-aza-2’dC incorporation into genomic DNA and the potency of decitabine in inhibiting cancer cell growth was highly correlated to those of azacitidine (Supplementary Figure 14A, 15A). Since decitabine cannot be incorporated into RNA, this high correlation provides further support that azacitidine exerts growth inhibition through the fraction that gets incorporated into DNA. In contrast to 5-aza-2’dCTP, gemcitabine or cytarabine triphosphate are not substrates for DCTPP1 (Carson et al, 2011; Requena et al, 2016). Consistent with this substrate selectivity, there was no appreciable synergy for the combination of triptolide and gemcitabine or cytarabine in multiple cell lines (Supplementary Figure 16A-C).

## DISCUSSION

Although nucleoside analog DNMTi have been effective for treatment of MDS and leukemia, they have shown limited single-agent clinical efficacy for the majority of human cancer types. Using a chemical screen of known drugs with subsequent validation and mechanistic studies, we identified that triptolide, a major active ingredient of *Tripterygium wilfordii*, and several of its analogs showed strong synergy with nucleoside analog DNMTi in suppressing growth and survival of cancer cells *in vitro* and enhanced efficacy of inhibiting xenograft tumor growth at well-tolerated doses *in vivo*. We used chemical biology, genetic approaches (knockout and overexpression of XPB or DCTPP1) and structural analysis to determine that the mechanism of synergy between triptolide and decitabine was attributable to targeting DCTPP1. By inhibiting DCTPP1-mediated hydrolysis of 5-aza-2’-deoxycytidine triphosphate, triptolide and its analogs i) enhanced the genomic incorporation of nucleoside analog DNMTi, ii) led to trapping and degradation of DNMT1, and to some extent DNMT3A and DNMT3B, and iii) maintained or even enhanced DNA demethylation and decitabine induced gene re-expression programs despite the synergistic cancer selective cytotoxicity of the combination.

Consistent with this notion, decitabine incorporation into genomic DNA and its efficacy were significantly associated with the level of expression of DCTPP1, but not XPB or any other enzymes involved in metabolism of deoxycytidine or decitabine en route to incorporation into genomic DNA in a series of cancer cell lines (Figure 8A, 8B, 8C). Knocking out DCTPP1 in cancer cells with intermediate to high DCTPP1 expression sensitized these cancer cells to single agent decitabine, while overexpressing DCTPP1 in cancer cells with low to intermediate DCTPP1 expression conferred resistance to single agent decitabine. These data suggest that DCTPP1 may represent a major cancer cell intrinsic mechanism of primary resistance to nucleoside analog DNMTi. Interestingly, a recent study has shown that DCTPP1 nuclear expression is often significantly enhanced in the majority of solid organ cancers studied compared to normal counterparts (Zhang et al., 2013), consistent with the poor activity of nucleoside analog DNMTi as single agents in solid organ cancers. This notion is further supported by our analysis of TCGA RNA-seq data showing that the majority of major solid organ cancers exhibited significantly higher DCTPP1 expression compared to that in hematological malignancies. (Figure 8G). Other enzymes such as cytidine deaminase, deoxycytidine kinase, and the nucleotide transporters hENT1/2, may also be involved in cell intrinsic and extrinsic resistance mechanisms as has been suggested by prior studies (Ebrahem et al., 2012; Kroep et al., 2002; Mahfouz et al., 2013; Qin et al., 2011; Stresemann and Lyko, 2008; Wu et al., 2015; Zauri et al., 2015), but their level of mRNA expression does not appear to be rate limiting in determining the degree to which decitabine can get incorporated into genomic DNA.

Inhibition of DCTPP1 also sensitized to the nucleoside DNMTi azacitidine, but did not sensitize to other cytidine analog drugs such as gemcitabine and cytarabine. This is likely due to the lack of enzymatic activity of DCTPP1 for the triphosphate metabolites of gemcitabine and cytarabine (Corson et al., 2011; Requena et al., 2014) Targeting DCTPP1 may also have some utility as a single agent by inhibiting the sanitation of damaged deoxycytidine nucleoside pools and by promoting random salvage incorporation of various naturally occurring modified cytidine nucleosides, such as 5-methylcytosine, hydroxymethylcytosine, formylcytosine, thereby interfering with their tightly regulated signaling functions in cancer cells (Corson et al., 2011; Gad et al., 2014; Requena et al., 2014; Song et al., 2015; Zauri et al., 2015). DCTPP1 may thus play a role in safe-guarding cancer cells against genomic and epigenomic instability.

The findings here provide strong rationale for advancing a combination of triptolide or its analogs with existing nucleoside analog DNMTi for testing in clinical trials. There is mounting interest in using nucleoside analog DNMTi alone or in combination with other epigenetic drugs and to use these combinations to prime sensitivity to subsequent or concomitant administration of other agents including targeted drugs, chemotherapeutic agents, and immunotherapies. In our *in vitro* treated cell model, there were significant enhancement of global demethylation and gene expression profiles, especially in interferon-α response genes and interferon-γ response genes (Figure 7D-F). Given the strong efficacy and safety and enhanced epigenetic effects seen in our *in vitro* and *in vivo* models for the combination of decitabine with triptolide and its DCTPP1-inhibiting analogs, these combinations may be more effective than DNMTi monotherapy when deployed in such trials. In the future, development and testing of selective DCTPP1 inhibitors, such as compounds with diverse heterocyclic scaffolds that have been recently reported, in combination with nucleoside analog DNMTi would also warrant clinical testing (Llona-Minguez et al., 2017a; Llona-Minguez et al., 2017b; Llona-Minguez et al., 2016). Based on our mechanistic studies, we can evaluate the utility of DCTPP1 gene and protein expression and the degree of decitabine genomic incorporation as predictive and pharmacodynamic biomarkers to inform such clinical trials. Thus the combination of triptolide and its DCTPP1 inhibiting analogs with nucleoside analog DNMTi is poised for near-term clinical translation. Finally, in future clinical studies, we can evaluate the potential of DCTPP1 expression and/or activity levels as a predictive biomarker for sensitivity to nucleoside-analog DNMTi monotherapy in MDS/leukemia.

## EXPERIMENTAL PROCEDURES

### Materials and cell culture

All the cell lines, unless otherwise stated, were obtained from American Type Culture Collection (ATCC) and grew in the recommended medium. Human prostate epithelial cell line RWPE1 was purchased from Lonza (Basel, Switzerland) and were cultured in PrEGM medium (Lonza). All cell lines were maintained in a humidified incubator with 5% CO_2_ at 37°C. All chemicals, unless otherwise stated, were purchased from Sigma Aldrich (St. Louis, MO) and had purity greater than 98%. [α -^32^P]-dCTP was purchased from Perkin Elmer (Waltham,MS); 5-methyl-2’-deoxycytidine (5mC), 5-azacitidine-^15^N_4_ (5AC-^15^N_4_), 2’-deoxycytidine-^13^C^15^N_2_ (2dC-^13^C^15^N_2_), and 5-methyl-2’-deoxycytidine-d3 (5mC-d3) were purchased from Toronto Research Chemical (Toronto, ON).

### Clinical drug library screen

3800 clinical compounds were included in Johns Hopkins Drug Library (JHDL) (Chong et al., 2006; Kamiyama et al., 2013; Platz et al., 2011). Human prostate cancer DU145 cells were treated with 100 nM Decitabine (DAC) or DMSO for 4 days in T75 flasks. Cells were then split and seeded into 96-well plates at 5,000 cells/well in complete media and allowed to adhere overnight. On day 5, JHDL drugs were added to the plates in duplicate wells at a final concentration of 5 μM for a total of 72 hours, with 16 wells left untreated as no-drug controls. After the first 48 hours in JHDL compounds, 1 μCi [^3^H]-thymidine was added, and incubated for an additional 24 hours (in both drug and [^3^H]-thymidine). Cells were then trypsinized and transferred onto FilterMat-A glass fiber filters (Wallac, Turku,Finland) using the Harvester-96 cell harvester (Tomtec, Hamden, CT). After washing with distilled water five times to remove free [^3^H]-thymidine, the incorporated [^3^H]-thymidine was read on a MicroBeta plate reader (Perkin Elmer,MA,USA), and the mean percentage of inhibition was calculated relative to no-drug control wells. In each group (DAC or DMSO pretreated cells), JHDL compounds demonstrating greater than 50% inhibition of [^3^H]-thymidine incorporation considered primary hits. After filtering out those compounds that are used only topically, 66 compounds representing a diverse array of classes were selected for combination titration studies across a wide dose range of each hit and of decitabine (0 nM to 10 μM each drug). For synthetic lethality and synergy analysis, the Combination Index (CI) was calculated with CompuSyn software using Chou-Talalay method (CI < 1 represents synergy) (Chou, 2010)and sum of Bliss independence index was calculated with Graphpad Prism 5 (Bliss independence index > 0 represents synergy) (Wei et al., 2012).

### *In vitro* DCTPP1 pyrophosphatase activity assays

[α -^32^P]-dCTP was used to detect dCTP hydrolysis by DCTPP1 with a thin layer chromatography assay. Briefly, a 20 μL reaction mixture contained 50 mM HEPES(pH=8.0), 100mM NaCl, 5 mM MgCl_2_, 1mM Tris (2-carboxyethyl) phosphine hydrochloride (TCEP), indicated concentration of triptolide analogs, 10 μM dCTP, 1 μCi [α -^32^P]dCTP, 100 μg/ml BSA, 10nM His-DCTPP1. The reactions were started by addition of DCTPP1 for 5min, and were stopped by addition of 5 μL of 0.5 M EDTA and dilution up to 100 μL with TE buffer. An aliquot of 1 μL reaction mixture was spotted on PEI-cellulose and the chromatogram was developed with 0.9 M guanidine hydrochloride (PH=6.0). The percent of dCTP hydrolysis to dCMP was quantified using a PhosphorImager (Molecular Dynamics, Sunnyvale, CA).

To enable measurement of DCTPP1 pyrophosphatase activity on multiple deoxycytidine triphosphate analog substrates, we measured the rate of DCTPP1 pyrophosphate production through a coupled ATP regeneration reaction followed by detection of luciferase-mediated luminescence. Briefly, the pyrophosphate generated by DCTPP1 after hydrolysis of dCTP or 5-aza-dCTP was converted by ATP sulfurylase to ATP, which was in turn used to produce luminescence by a luciferase assay. A 45 μL reaction mixture contained 50 mM HEPES (pH=8.0), 100mM NaCl, 5 mM MgCl_2_, 1mM TCEP, various concentrations of triptolide, 100 μg/ml BSA, 0.015U ATP sulfurylase, 10nM His-DCTPP1. The reactions were started by addition of various concentrations of 5-Aza-dCTP or dCTP, and incubated at room temperature at the indicated times followed by quenching via addition of 5 μL of 0.1 M EDTA. The luminescence produced by addition of 50 μL Buffer A (50mM HEPES(PH=8.0), 15 mM MgCl_2_, 5 μM Adenosine 5’-phosphosulfate (APS), 0.015 U luciferase, and 100 ug/ml D-luciferin, using injector mode of Microbeta 1450-023(Perkins Elmer). The amount of DCTPP1 mediated production of pyrophosphate from 5-aza-dCTP or dCTP was quantified comparing to a pyrophosphate standard curve. The Kcat and Km for each substrate was calculated by Prism Graphpad 5.0.

### *In vitro* XPB activity assays

The TFIIH complex containing XPB was purified and its DNA-dependent ATPase assay was performed as previously described (He et al., 2015; Titov et al., 2011). Briefly, a 10-μl reaction mixture contained 20 mM Tris (pH 7.9), 4 mM MgCl_2_, 1μM of ATP, 1 μCi [γ-^32^P]ATP (3000 Ci/mmol), 100 μg/ml BSA, 100nM AdMLP, 5nM TFIIH and indicated concentrations of triptolide or its analogs. The reactions were started by either addition of TFIIH for 2hr and stopped by addition of 2 μl of 0.5 M EDTA and dilution up to 100 μl with TE buffer. An aliquot of 1 μl reaction mixture was spotted on PEI-cellulose and the chromatogram was developed with 0.5 M LiCl and 1 M HCOOH. The percent of ATP hydrolysis was quantified using a PhosphorImager.

### Quantitation of genomic 5-aza-dC and 5-methyl-dC content by high-pressure liquid chromatography-tandem mass spectrometry (LC-MS/MS)

The 5-aza-deoxycytidine (5-aza-dC) and 5-methyl-deoxycytidine (5-methyl-dC) content in genomic DNA was determined by a high-pressure liquid chromatography/tandem mass spectrometry (LC-MS/MS) procedure as described previously, by the Johns Hopkins Analytical Pharmacology Core facility (Anders et al., 2016; Yegnasubramanian et al., 2008). Briefly, ∼2 to 5 μg genomic DNA in 50 μL of HPLC grade water was digested with 4 Units of Nuclease P1 (Sigma) at 65°C for 10 minutes in a digestion buffer containing 0.04 mM desferrioxamine mesylate (DFAM), 3.25 mM ammonium acetate pH 5.0, 0.5 mM zinc chloride in a final volume of 100 μL. Subsequently, 20 μL of 100 mM Trizma base, pH 8.5 was added and this reaction was treated with 4 Units of Alkaline Phosphatase (Roche Life Science) at 37°C for 1 hour. Following incubation, 20 μL of 300 mM ammonium acetate, pH 5.0 and 6 μL of 0.25 mM DFAM in 50nM EDTA was added to stop digestion. Standards and quality control (QCs) samples were prepared by adding known concentrations of decitabine, 2′-deoxycytidine (2dC), and 5-methyl-2′-deoxycytidine (5mC) into blank digest matrix (all solvents used during DNA digest except for the enzymes). All samples, standards and QCs were prepared for analysis by adding 20 µL internal standard (5-azacitidine (5AC)-^15^N_4_ 2dC-^13^C^15^N_2_, and 5mC-d3 in water) to 100 µL of sample and vortex-mix briefly. The samples were injected onto a LC-MS/MS system consisting of a Waters Acquity UPLC (Milford, MA) interfaced with an AB Sciex 5500 triple quadruple mass spectrometer (Foster City, CA). Chromatographic separation was achieved using Thermo Hypercarb porous graphite analytical column (100 x 2.1 mm, 5 µm, Waltham, MA) running isocratic elution with a mobile phase consisting of 10 mM ammonium acetate:acetonitrile with 0.1% formic acid (70:30, v/v) at a flow rate of 0.3 mL/minute. The mass spectrometer was run in positive electrospray ionization mode monitoring for the following MRM transitions: 5-aza-dC (DAC): 228.9 ➔ 113.0, 2dC: 228.0 ➔112.0, 5-methyl-dC: 242.0➔126.0, 5AC-^15^N_4_ (internal standard): 249.0 ➔ 117.0, 2dC-^13^C^15^N_2_ (internal standard): 230.8 ➔ 115.0, and 5mC-d3 (internal standard): 245.8 ➔129.0. The calibration range was 2 – 400 ng/mL for 5-aza-dC, 5-1000 ng/mL for 5-methyl-dC, and 50-10,000 ng/mL for 2dC. All analytes used quadratic regression with 1/x^2^ weighing. Results were reported as 5-aza-dC (DAC) and 5mC content per thousand 2dC.

### Western blot detection of DNMT, DCTPP1and XPB proteins

Cells were collected by trypsinization, washed with PBS, and lysed in RIPA buffer (ThermoFisher) supplemented with protease and proteosomal inhibitors. Lysates were denatured in NuPage LDS Sample buffer 4X (Invitrogen, US) at 100°C and 15 μg of protein were separated with 4–12% Bis-Tris gels (Invitrogen, US) and transferred to polyvinylidene fluoride (PVDF) membrane using NUPAGE transfer buffer (Invitrogen, US). Membranes were incubated with Odyssey blocking buffer (Li-Cor, Lincoln, NE) prior to incubation with appropriate antibodies (anti-DNMT-1,-3A,-3B antibody, 1:1000, Sigma; anti-DCTPP1 antibody, 1:500, Abcam; anti-XPB antibody, 1:500, Abcam; anti-β-actin antibody, 1:5000 Cell Signaling) overnight at 4°C. After three 5 minute washes with PBS-Tween20 (0.05%), goat anti-rabbit or goat anti-mouse secondary antibody (1:5000, LI-COR) was applied for 1 hour at room temperature, followed by three washes with PBS-Tween20 (0.05%). Visualization and quantification was carried out with the LI-COR Odyssey scanner and software (Li-Cor, Lincoln, NE).

### Clonogenic survival assay

Clonogenic assays were performed to assess long-term cell survival. Human prostate cancer cell lines, DU145 and PC3 cells were plated in duplicates in a density of 2,000 cells/well in 6-well plates for adhering overnight. Decitabine was freshly dissolved in DMSO and diluted with PBS to indicated concentration and added to culture plates within 15 minutes. Cells were cultured for 10-14 days to allow formation of colonies. After 30 minutes fixation with 20% methanol, the clones were stained with crystal violet (Sigma, US) and the colony areas were calculated with ImageJ (Guzman et al., 2014). All dishes from one cell line were stained at the same time point. The average colony area for each condition was normalized to that in DMSO-treated controls for each cell line and each separate experiment. Statistical significance of differences was assessed by Student’s *t*-tests.

### Alamar Blue cell viability assay

Cells were seeded into 96 well plates one day before compound treatments. After incubating with appropriate drugs for 96 hours, 20 µL Alamar Blue reagent (Thermo Fisher, USA) was added into each well and incubated for 2 hours. The fluorescence intensity was read on a microplate reader at 590nm wavelength.

### Growth curve assay

500 cancer cells were seeded per well in 48 well plates one day before compound treatments. Decitabine, triptolide and other drugs were freshly prepared and added to culture plates at indicated concentrations. For growth curve experiments, growth data were determined by imaging cell confluence at 6 hour intervals at each of 16 fields per well using the IncuCyte™ ZOOM System (Essen Bioscience, MA) in the Johns Hopkins SKCCC Imaging Core Facility.. The percent confluence was calculated with Incucyte built-in software. Growth curves were analyzed and constructed with Graphpad Prism software 5.0 (La Jolla, CA).

### Growth inhibition of subcutaneous xenografted tumors

Protocols for all animal experiments conducted at Johns Hopkins University were approved by the University Animal Care and Use Committee. Human prostate cancer DU145 xenografts were generated by subcutaneous injection of 2×10^6^ cells into 8-10 week old athymic nude mice (Harlan, Indianapolis, IN). Tumor volumes (calculated as length x (width)^2^/2) were measured every week. When the tumors reached approximately 200 mm^3^ in size, 10 mice were randomly divided into each treatment arm and received intraperitoneal injection of 100ul of appropriate drug as outlined in Figure 3A: control (normal saline, Arm 1), decitabine (three 1 mg/kg doses given i.p. every other day on week 1 of two 3-week cycles, Arm 2), triptolide (0.2 mg/kg doses given each day for 5 days with two days’ rest each week for two 3-week cycles, Arm 3) or both (Arm 4).

For analysis of systemic toxicity, mouse body weight was measured every week; whole blood samples were collected from three randomly selected mice in each group at baseline, at week 2, and at completion of treatment (week6) by submandibular bleeding. Mouse blood samples were processed within four hours for measurement of complete blood count/hematology and clinical chemistry parameters by the Johns Hopkins Phenotyping Core facility.

### Lentiviral-mediated stable gene knockout

pLenti-CRISPR.V2 plasmids containing DCTPP1 CRISPR guide RNA GCTTACATGTCCTCGAGCG), or XPB CRISPR guide RNA (AAACGGGCGTGCACGTTCGG), or non-targeting control were obtained from Genscript (Piscataway, NJ). Packaged viral particles were used to infect selected cancer cell lines using polybrene media (at a final concentration of 8 μg/ml). Transduced cells were selected by puromycin (1-2.5 μg/ml) (Sigma). Knockdown efficacy was determined by western blotting and cells were used for further studies as described.

### Lentiviral-mediated stable gene overexpression

pLenti-6 containing DCTPP1 or XPB ORF were shared from Dr. Jun O Liu’s lab in Johns Hopkins University. Packaged viral particles were used to infect selected cancer cell lines using polybrene media (at a final concentration of 8 μg/ml). Transduced cells were selected by blasticidin (5-10 μg/ml) (Sigma). Expression efficacy was determined by western blotting and cells were used for further studies as described.

### X-ray Crystallography

#### Constructs and Cloning

The gene for full length human DCTPP1 was codon optimized for bacterial expression and synthesized by Twist bioscience (www.twistbioscience.com). A slightly truncated crystallization construct comprising residues 21-130 (DCTPP1^(21-130)^) was designed using sequence alignments with existing crystal structures of mouse DCTPP1 (2A3Q, 2OIG). DCTPP1^(21-130)^ was generated by PCR and subcloned into a modified pET vector containing an N-terminal hexahistidine SUMO tag using ligation independent cloning (LIC) (Aslanidis and de Jong, 1990).

#### Expression and Purification

Expression of DCTPP1^(21-130)^ was conducted in BL21[DE]-RIL *E. coli* (www.agilent.com). Cells were grown in 2XTY at 37°C to OD_600_ of 0.5 before lowering the temperature to 18°C for 30 minutes, followed by induction with 300 μM IPTG. Cells were harvested after 16 hours at 18°C and resuspended in Buffer A (50 mM Tris-HCl (pH 7.8); 800 mM NaCl; 10% glycerol; 10 mM imidazole; 0.5 mM TCEP-HCl; 1 mM PMSF; 1 μg/mL Leupeptin; 1 μg/mL Pepstatin). Cells were lysed by sonication and clarified by centrifugation before passing the lysate over a 5 mL HisTrap-HP column (www.cytivalifesciences.com). The column was washed with 200 mL of Buffer A and eluted with 30 mL Buffer B (50 mM Tris-HCl (pH 7.8); 800 mM NaCl; 10% glycerol; 500 mM imidazole; 0.5 mM TCEP-HCl; 1 mM PMSF; 1 μg/mL Leupeptin; 1 μg/mL Pepstatin). SUMO-tagged DCTPP1^(21-130)^ was cleaved with SENP protease and dialyzed overnight against Buffer A. Cleaved DCTPP1^(21-130)^ was again applied to a 5 mL HisTrap-HP column and flow-through containing protein was collected and concentrated for application onto a size exclusion column. Concentrated protein was applied to a Sephacryl S-300 column (www.cytivalifesciences.com) pre-equilibrated in Buffer C (50 mM Tris 7.8; 500 mM NaCl; 10% glycerol; 0.5 mM TCEP). Peak fractions were pooled and concentrated to 20 mg/mL. Final glycerol concentration was brought to 30% for flash freezing and storage at −80°C.

#### Crystallization, Data Collection and Model Building

For crystallization, DCTPP1^(21-130)^ was dialyzed overnight against Buffer D (20 mM Tris 7.8; 500 mM NaCl; 0.5M TCEP). After dialysis, 96 μL of DCTPP1 (10mg/mL) was complexed with 4 μL triptolide (50mM in 100% DMSO) to a final concentration of 2 mM Triptolide. The DCTPP1^(21-130)^•triptolide complex was crystallized by hanging drop vapor diffusion. Protein and compound was mixed in a 1:1 ratio with crystallization buffer containing 100 mM Tris-HCl (pH 7.5), 8% PEG3350, and 300 mM proline and allowed to equilibrate by vapor diffusion at 20°C. Crystals were cryoprotected in crystallization buffer supplemented with 25% glycerol and 2 mM triptolide before plunge freezing in liquid nitrogen. Diffraction data were collected at NSLS-II beamline 17-ID-1 (AMX) and auto-processed using FASTDP (Winter and McAuley, 2011). A molecular replacement solution was obtained by using 2OIG as a search model, stripped of ligands and waters using PHASER (McCoy et al., 2007) in the PHENIX software suite (Adams et al., 2010). Model building was conducted using COOT (Emsley and Cowtan, 2004). Refinement was carried out in PHENIX (Adams et al., 2010) and figures were generated using PYMOL (www.pymol.org). Coordinates have been deposited in the PDB (accession number 7MU5).

### RNAseq

DU145 cells were exposed to either 2 nM triptolide alone, 33 nM decitabine alone, or both 2 nM triptolide and 3 nM decitabine once daily for 3 days, followed by 17 more days of treatment-free growth. Following this 20-day period, RNA was extracted and purified for RNA-seq using Qiagen Allprep method (Germantown, MD). Complementary DNA libraries were prepared with Illumina’s TruSeq Stranded Total RNA Sample prep and were sequenced on an Illumina HiSeq System. Reads were aligned to the transcriptome using STAR, quantified using a gene transfer format (GTF) file containing gene models, with each model representing the structure of transcripts produced by a given gene, and normalized to transcripts per million (TPM). The significance and magnitude of differential expression between treatment groups was determined using the R package *DEseq2* (Love et al., 2014). Heatmaps were generated using the CRAN package *gplots*. The Wilcoxon rank sum test was performed on the log_2_ fold change (DAC+TPL versus DAC) to assess whether genes upregulated by decitabine alone were enriched for further upregulation by combination treatment of decitabine and triptolide. Gene set enrichment for increased expression with treatment of DAC+TPL versus DAC was performed with GSEA against HALLMARK gene sets containing between 15 and 500 genes. HALLMARK gene sets were downloaded from gseaftp.broadinstitute.org/.

### Global Genomic DNA methylation assays

Cells were treated with vehicle control, decitabine, triptolide or in combination for 5 days. Genomic DNA was extracted with DNeasy kit (Qiangen, Germantown, MD) according manufacturer’s protocol. The global DNA methylation were analyzed by direct Pyrosequencing after bisulfite conversion of DNA in EpigenDX (Hopkinton, MA).

### Global gene expression and methylation analysis

Gene expression profiles for queried cells were analyzed using Agilent Human 44K expression arrays (Agilent Techonologies, Santa Clara, CA). Data was loess-normalized to correct for dye and ratio bias at different signal intensities. Global methylation analysis was performed using the Illumina Infinium Human Methylation27 BeadChip (Illumina, Inc. San Diego, CA). Data quality was verified using *in vitro* methylated DNA (IVD) and DNMT1 and 3b double knock-out HCT-116 cells (DKO), which is highly depleted of methylation. Data from the above two platforms were analyzed with R and BioConductor.

### Statistics

Statistical analyses were performed using Prism GraphPad v 5.0 software (San Diego, CA), unless described separately above. The *a priori* level of significance was set at 0.05.

## AUTHOR CONTRIBUTIONS

J.L., J.O.L., W.G.N. and S.Y. designed the study, analyzed data, and wrote the article. J.L., A.R., P.N. and J.S.S. performed Johns Hopkins Drug Library screen and synergy analysis. J.L., W.G.N. and S.Y. designed and analyzed animal studies for human prostate cancers. N.C., A.M.V. and J.L. conducted all animal studies. J.L., J.Y.Z., N.M.A., T.W., H.G., M.A.R. and S.Y. designed and conducted all the experiments related to LC-MS/MS. J.L., D.E., J.Y.Z. and A.M.V. conducted all the *in vitro* cell viability assay, cell growth assay and clonogenic survival assays. Q.L.H., J.L., J.O.L., W.G.N. and S.Y. designed and conducted all the *in vitro* DCTPP1 and XPB assays. J.L., R.C.S., R.V.C., W.G.N. and S.Y. designed and conducted all the global gene expression and methylation experiments. G.H., J.M.B., designed and conducted all the X-ray crystallography. All authors contributed to the writing, editing, and approval of the final manuscript.

## ACKNOWLEDGMENTS

Funding:

This work was supported by NIH/NCI grants P50CA058236, R01CA183965, R01CA070196, P30CA006973, S10RR026824, UL1TR003098, Prostate Cancer Foundation Challenge Awards, Patrick C. Walsh Koch Research Award, the Ellen B. Masenheimer Fellowship (S.Y.), the Flight Attendant Medical Research Institute (J.O.L.) and the Commonwealth Foundation. This research used beamlines AMX(17ID-1) and FMX(17ID-2) of the National Synchrotron Light Source II (NSLS-II), a U.S. Department of Energy (DOE) Office of Science User Facility operated for the DOE Office of Science by Brookhaven National Laboratory under Contract No. DE-SC0012704. The Center for BioMolecular Structure (CBMS) at NSLS-II is primarily supported by the National Institutes of Health, National Institute of General Medical Sciences (NIGMS) through a Center Core P30 Grant (P30GM133893), and by the DOE Office of Biological and Environmental Research (KP1605010).

