## Supplemental figures legends for "Triptolide sensitizes cancer cells to nucleoside DNA methyltransferase inhibitors through inhibition of DCTPP1-mediated cell-intrinsic resistance"

**A**

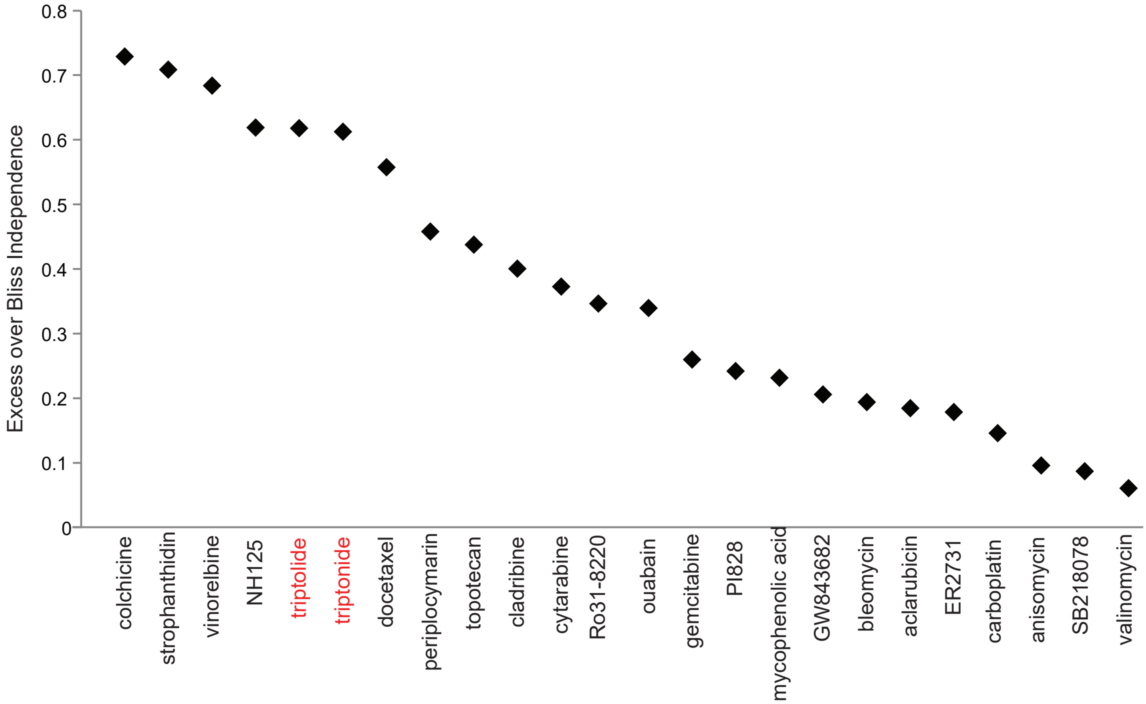


**Figure S1. (Related to Figure 1.) Scatter plot of 24 synergistic compounds with decitabine ranked by sum of excess over Bliss Independence as the synergy score.** Triptolide and triptonide are specified in red color. Values greater than zero indicate increasing levels of synergy.

A

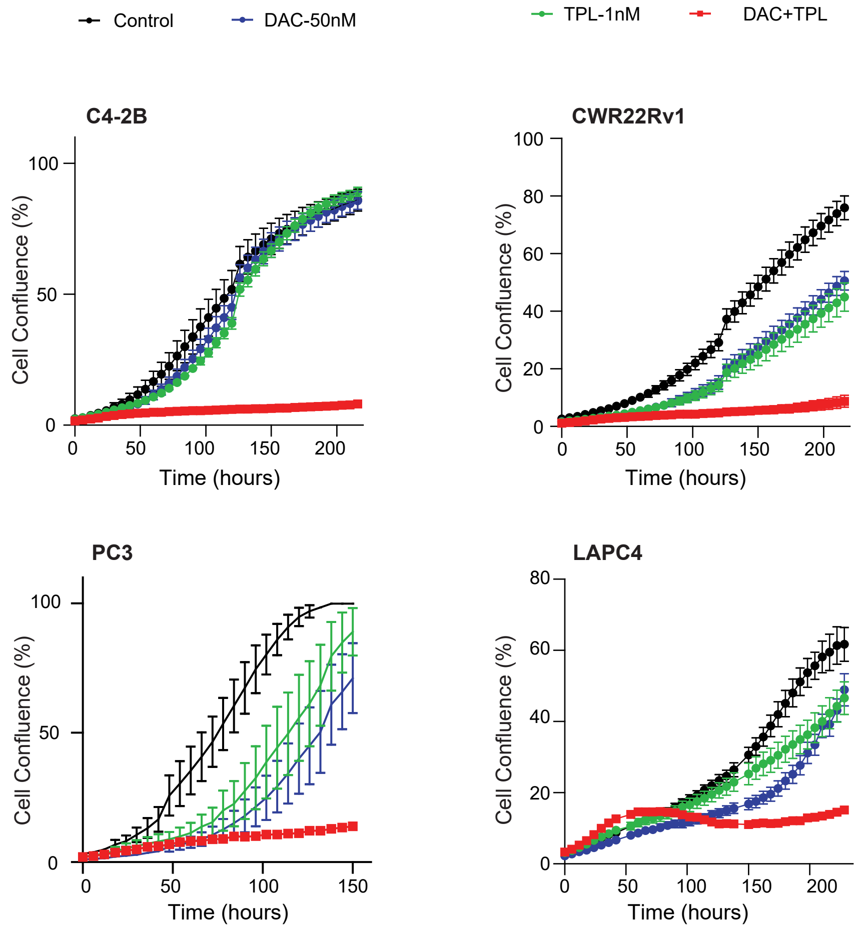


**Figure S2. (Related to Figure 2.) Triptolide synergistically sensitizes cancer cells to nucleoside analog DNMT inhibitors *in vitro*.** Inhibition of growth of human prostate cancer cell lines C4-2B, CWR22Rv1, LAPC4 and PC3 treated with vehicle control, 50nM decitabine or 1 nM triptolide alone or in combination. Growth curves were measured as the percent of total confluence in each well over time. The mean percent confluence ± SEM for 16 fields in each condition are shown, with measurements taken every 6 hours throughout the course of the growth curve.

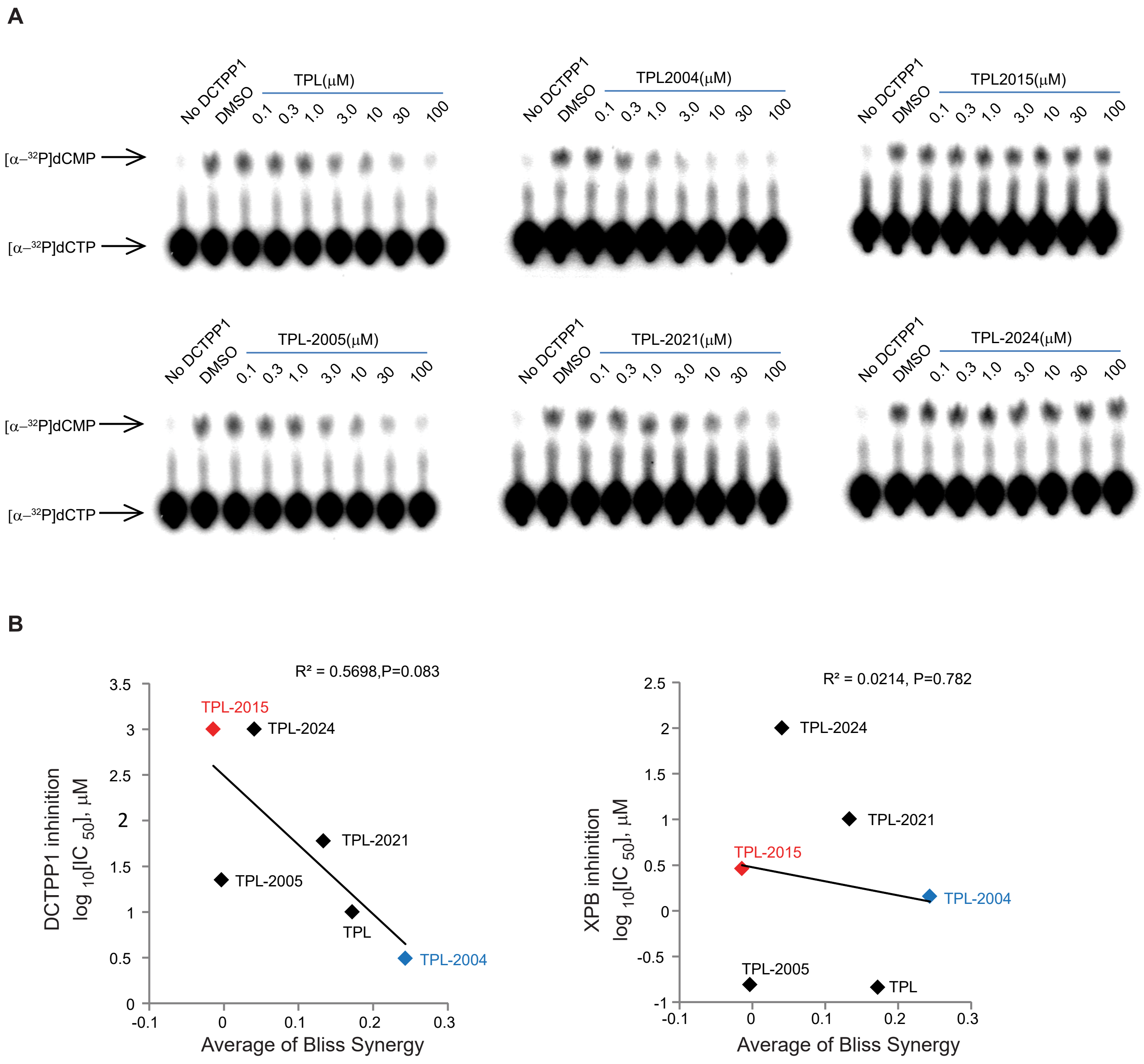


**Figure S3. (Related to Figure 4,Table1.) Inhibition of DCTPP1 activity by triptolide analogs and relationship to synergy with decitabine.** (A). Thin layer chromatography DCTPP1 pyrophosphatase activity assay using 1 μCi [α -$^{32}$P]dCTP, with a dose response of triptolide and its analogs. The percent of dCTP hydrolysis to dCMP was quantified using PhosphorImager. Shown are typical images of duplicate reactions. (B).Correlation between synergy of decitabine and triptolide analogs (measured as average of Bliss synergy across all doses) in inhibiting DU145 cell growth (measured by growth curve analysis as in Fig 4D) and the potency of inhibiting DCTPP1 (left panel) or XPB (right panel) activity *in vitro* for a series of triptolide analogs. The synergy between decitabine and triptolide analogs was more correlated with the potency of triptolide analogs in inhibiting DCTPP1 activity than in inhibiting XPB.

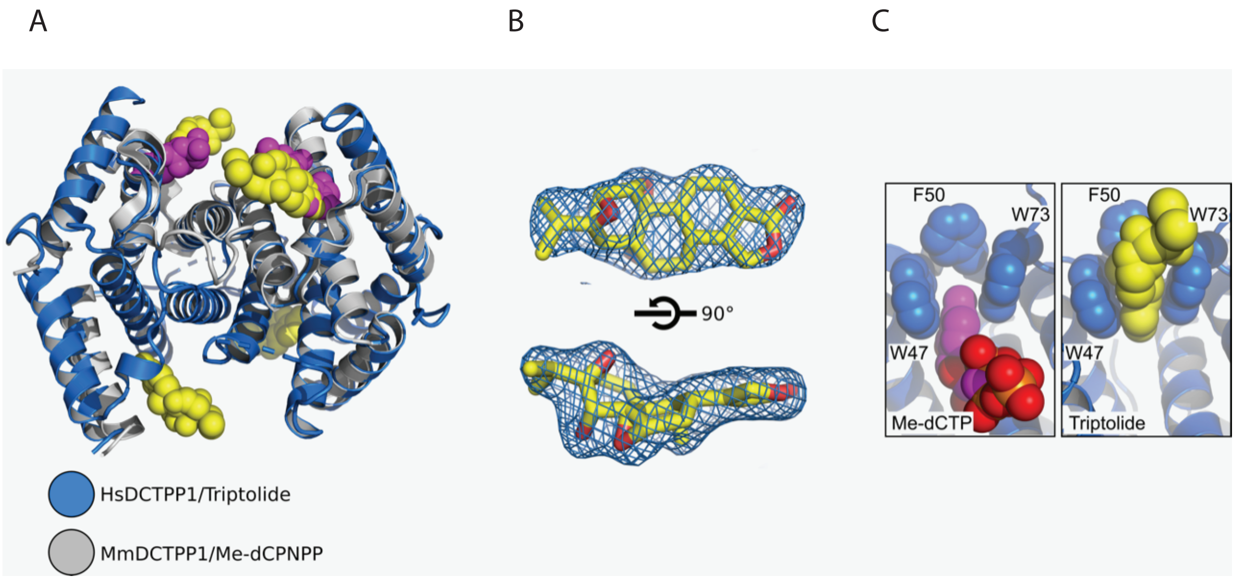


**Figure S4. (Related to Figure 4.) Superposition of DCTPP1 binding with triptolide and Me-dCPNPP**. (A) Structural alignment of the structure of triptolide bound to human DCTPP1$^{(21\text{-}130)}$ (blue ribbon) and mouse DCTPP1 bound to Me-dCPNPP (grey ribbon – PDB# 6SQZ). Triptolide and Me-dCPNPP are illustrated as yellow and magenta spheres, respectively. (B) Electron density for a 2Fo-Fc simulated annealing omit map contoured at $1\sigma$ for triptolide bound to DCTPP1. (C) Space filling representation of the hydrophobic binding pocket shared by Me-dCTP and triptolide. Me-dCTP is illustrated as magenta spheres with phosphorous and oxygen atoms colored orange and red, respectively. Triptolide is illustrated as yellow spheres as in (A).

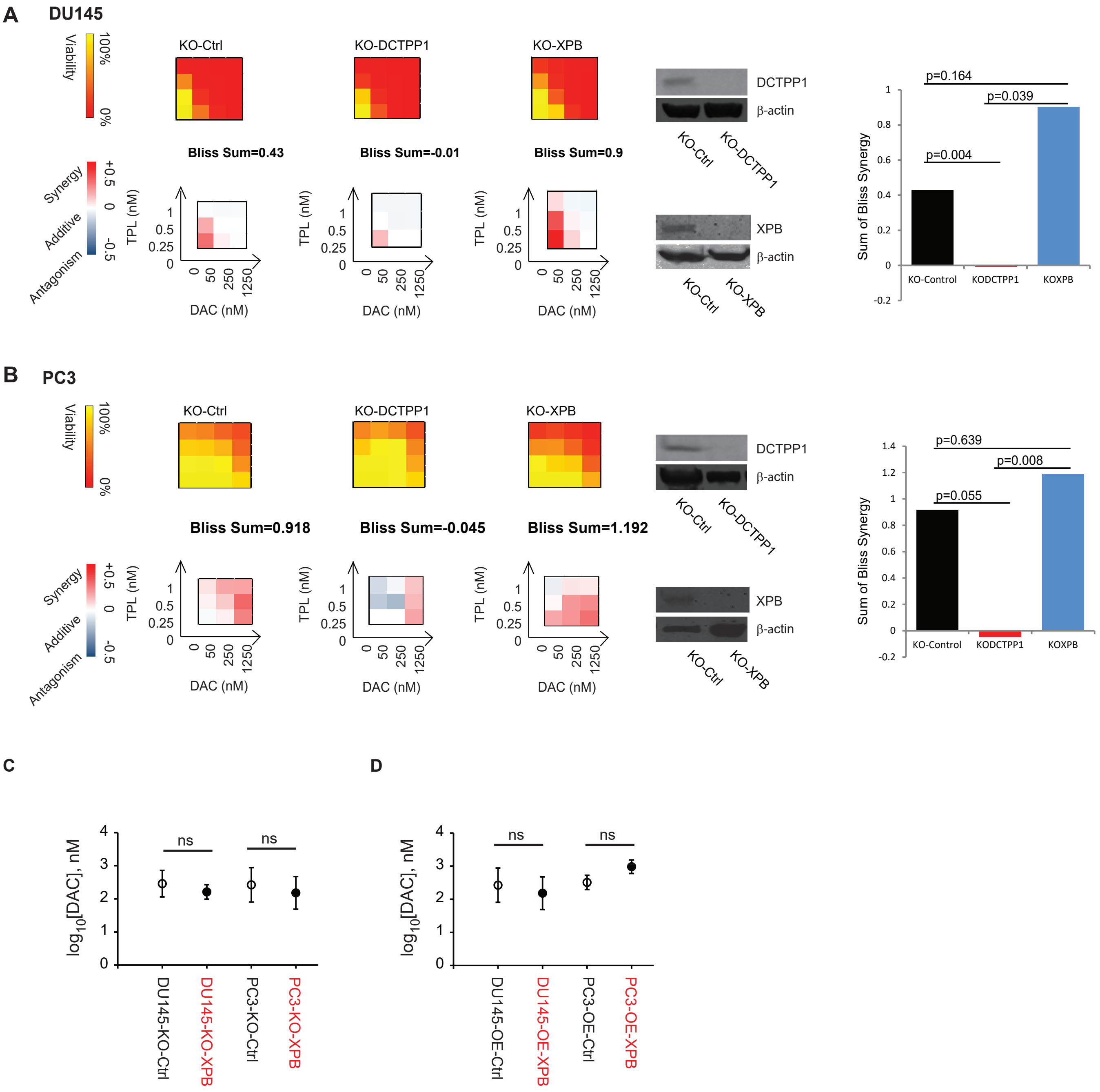


**Figure S5. (Related to Figure 5.) The synergy of triptolide and decitabine is dependent on DCTPP1.** Human prostate cancer cell lines DU145 (A) and PC3 (B) were transduced with CRISPR-Cas9 with non-targeting controls (KO-Control), or DCTPP1 guide-RNA to knockout DCTPP1 (KO-DCTPP1) or XPB guide-RNA to knockout XPB (KO-XPB) confirmed by immunoblotting. Inhibition of cell viability and the degree of synergy of combinations of decitabine and triptolide in indicated dose series of the transduced isogenic cell lines were developed as in Figure 7. Knocking out XPB in DU145 and PC3 cells had no significant change of overall synergy between triptolide and decitabine (histograms, right panel) compared to isogenic control cells. Knocking out DCTPP1 in both DU145 and PC3 cells completely abolished overall synergy (Bliss Sum<0). (C, D). Data represent $\log_{10}$ [IC50, DAC (nM)] and 95% confidence intervals of the XPB knockout or overexpression cell lines. NS, non-significant.

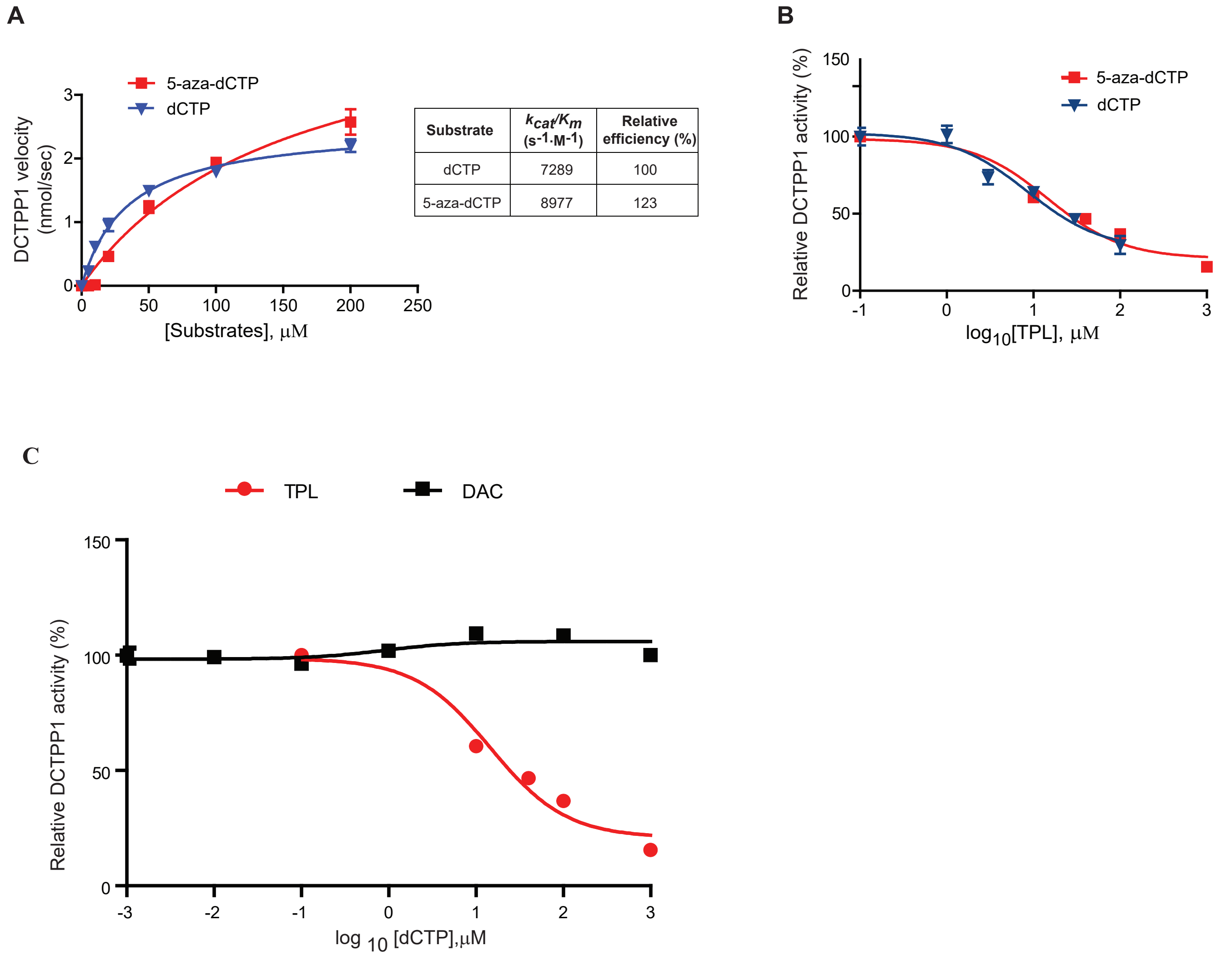


| Substrate | $k_{cat}/K_m$ ($s^{-1} \cdot M^{-1}$) | Relative efficiency (%) |
|---|---|---|
| dCTP | 7289 | 100 |
| 5-aza-dCTP | 8977 | 123 |

**Figure S6. (Related to Figure 6.) dCTP and 5-aza-dCTP are optimal substrates in *in vitro* DCTPP1 assay.** (A). Hill plot of DCTPP1 enzymatic pyrophosphatase activity on a range of dCTP and 5-aza-dCTP substrate concentrations. Each value represents the mean ± SEM of triplicate measurements of the initial velocity of DCTPP1 pyrophosphatase activity for each substrate. (B).Triptolide inhibits DCTPP1 pyrophosphatase activity, measured via a coupled luminescence assay (see materials and methods), on dCTP and 5-aza-dCTP across a dose response. Shown are the mean ± SEM of triplicate measurements. (C). Decitabine failed to inhibit DCTPP1 across a wide dose range as indicated. Triptolide served as a positive control for inhibition of DCTPP1 activity. Each value represents the mean ± SEM of triplicate treatments.

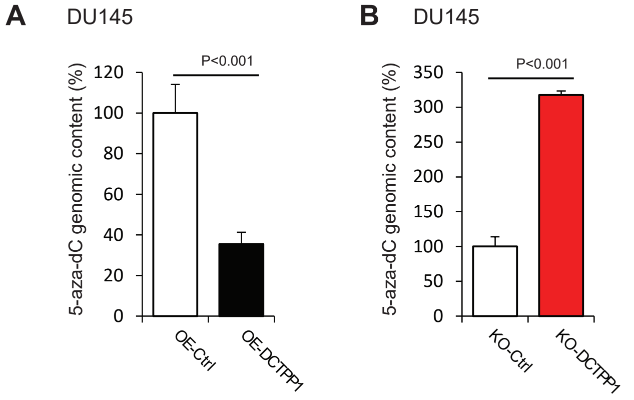


**Figure S7. (Related to Figure 6.) The incorporation of decitabine into genomic DNA is strongly influenced by modulating DCTPP1 levels.** (A). Overexpression of DCTPP1 in DU145 cells reduced decitabine incorporation into genomic DNA. 5-aza-2'-dC incorporation into genomic DNA, determined by LC-MS/MS, in DU145-OE-Control or DU145-OE-DCTPP1 cells treated with vehicle control or 1250nM decitabine for 2 days. The 5-aza-2'-dC content per 1000 2'-dC in DU145-OE-DCTPP1 cells was normalized to that of DU145-OE-Control cells. (B). Knockout DCTPP1 in DU145 cells enhanced decitabine incorporation into genomic DNA. 5-aza-2'-dC incorporation into genomic DNA, determined by LC-MS/MS, in DU145-KO-Control or DU145-KO-DCTPP1 cells treated with vehicle control or 1250nM decitabine for 2 days. The 5-aza-2'-dC content per 1000 2'-dC in DU145-KO-DCTPP1 cells was normalized to that of DU145-KO-Control cells. Shown in (A and B) are the mean ± SEM of triplicate treatments.

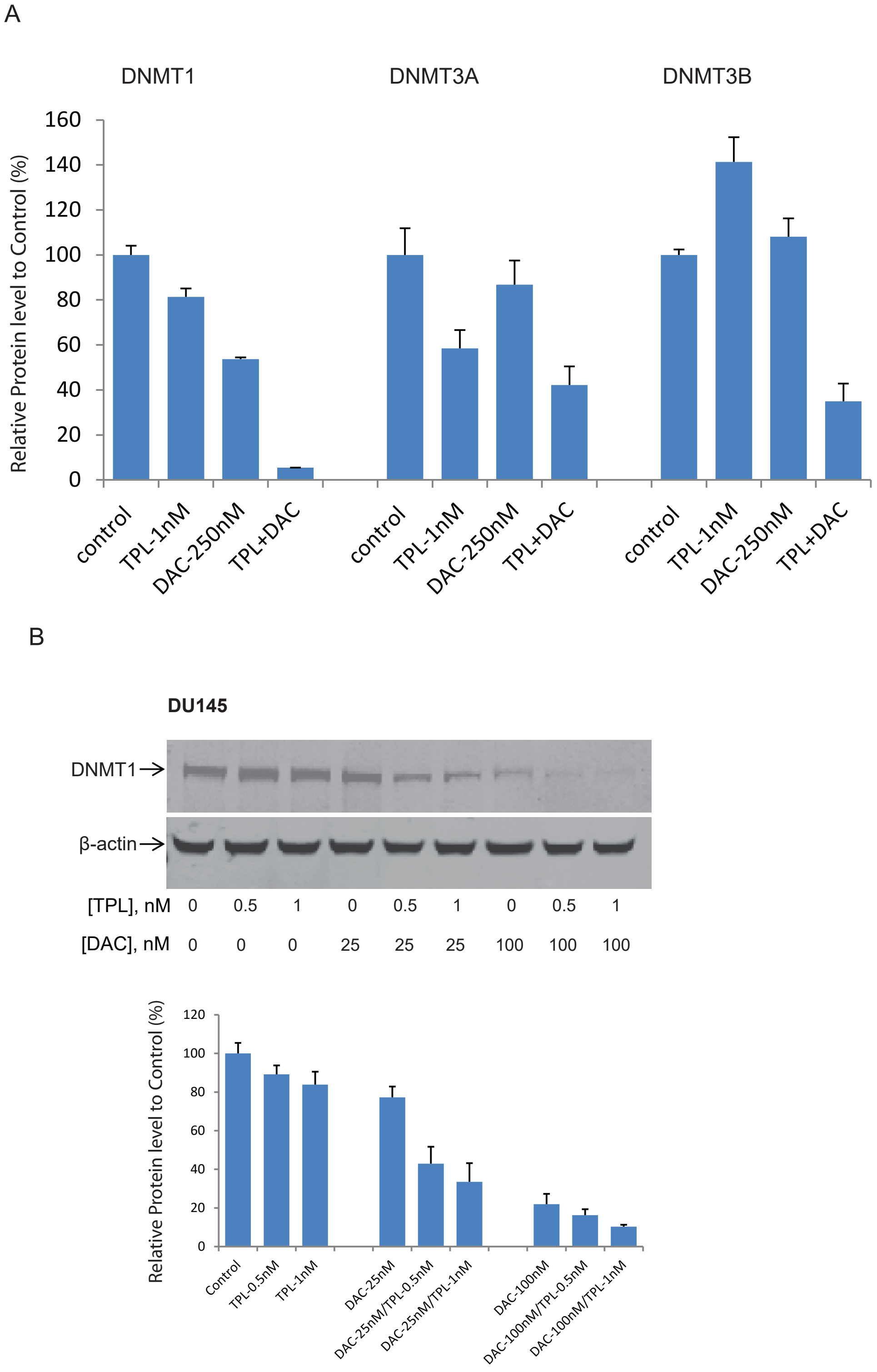


**Figure S8. (Related to Figure 6.) Triptolide treatment enhanced the decitabine induced DNMTs protein loss.** (A). Quantitation of immunoblots from Figure 6C of DNMT1,DNMT3a and DNMT3B levels in human prostate cancer DU145 cells treated with vehicle control, decitabine (250 nM) or triptolide (1 nM) alone or in combination for 24 hours. Densitometry was measured with Image-J software. (B). Western Blot of DNMT1 levels in human prostate cancer DU145 cells treated with a 2-dose response of each drug: vehicle control, decitabine (25nM, 100 nM) or triptolide (0.5 nM and 1 nM) alone or in combination for 24 hours. β-actin blotting is shown as protein loading control, and densitometry was measured as in (A).

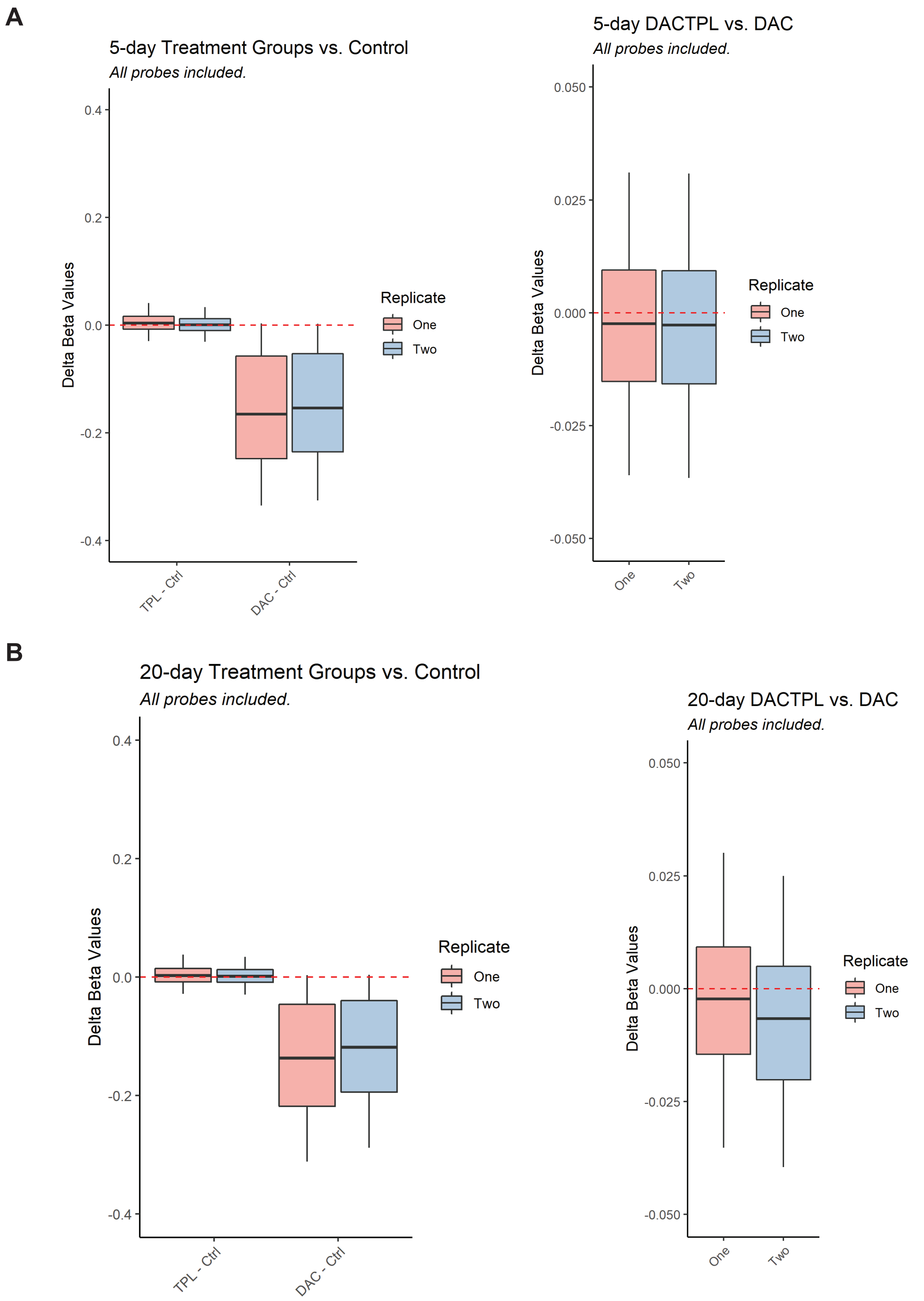


**Figure S9. Genome-wide methylation analyses in DU145 cells after treatment with triptolide and decitabine.** (A) Distribution plots of global DNA demethylation (delta beta values) across all CpG probes in Infinium platform in DU145 cells treated with vehicle control, 33 nM decitabine (DAC) alone or 2 nM triptolide (TPL) alone or in combination (DAC+TPL) for 5 days (A) or 20 days (B). The y-axis represents the difference in beta values at each probe between the two conditions indicated. Negative values indicate greater extent of demethylation in the first condition compared to the second. At both timepoints, the combination of decitabine and triptolide led to greater demethylation compared to decitabine alone ($P < 10^{-16}$, Wilcoxon signed rank test).

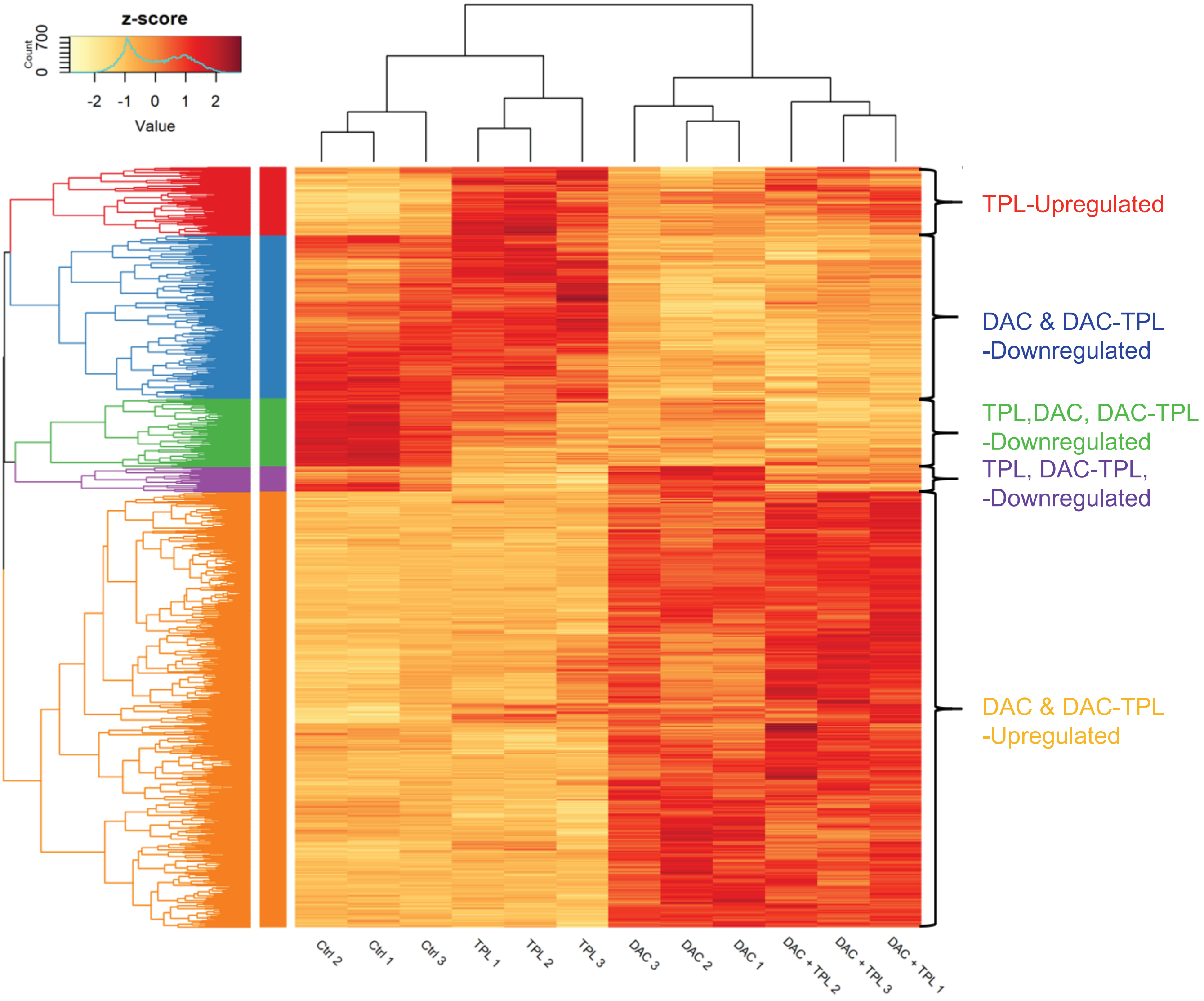


**Figure S10. (Related to Figure 7). The combination treatment of triptolide and decitabine changes gene expression profiles in DU145 cells.** Heatmap of DU145 expression profile change in RNA-seq with Vehicle control (Ctrl), 2nM Triptolide (TPL), 33nM Decitabine (DAC) or the combination (DAC+TPL). Shown are differentially expressed genes across conditions (n=3 each) at 20 days after treatment. Genes showing specific patterns of alteration across the conditions are clustered together and indicated.

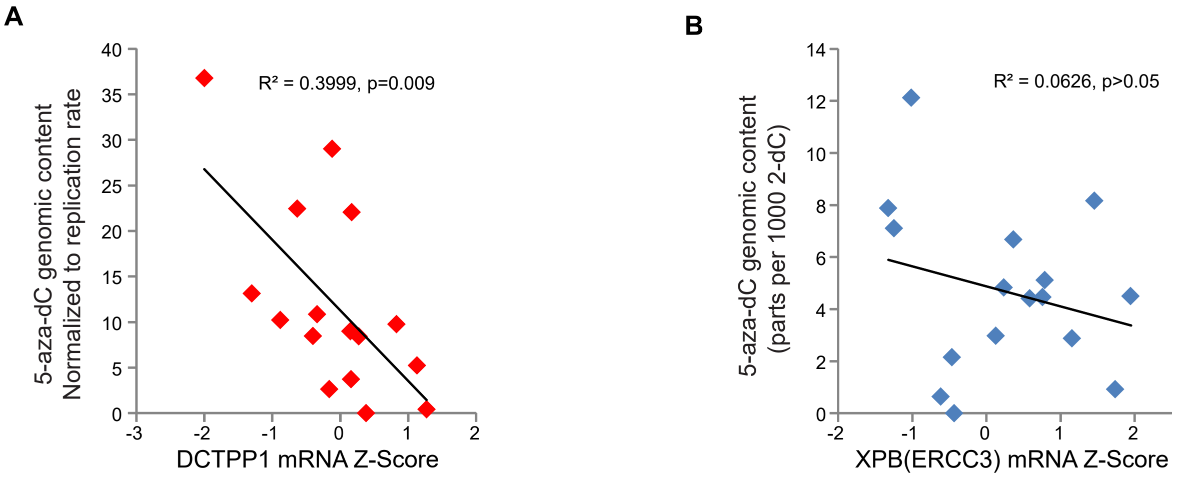


**Figure S11.(Related to Figure 8).DCTPP1 but not XPB expression levels are inversely correlated with 5-aza-dC incorporation.** (A) Correlation between levels of DCTPP1 mRNA expression and decitabine incorporation after normalization with cell proliferation activity among 16 cancer cell lines. Tritiated thymidine incorporation assays were carried out in parallel experiments with decitabine incorporation assays in order to normalize for differences in DNA replication rates among the different cell lines. The significant inverse correlation between decitabine incorporation rate and DCTPP1 mRNA expression level is preserved after normalization to tritiated thymidine incorporation. (B).There is no significant correlation between XPB mRNA expression levels with decitabine incorporation levels among 16 cancer cell lines.

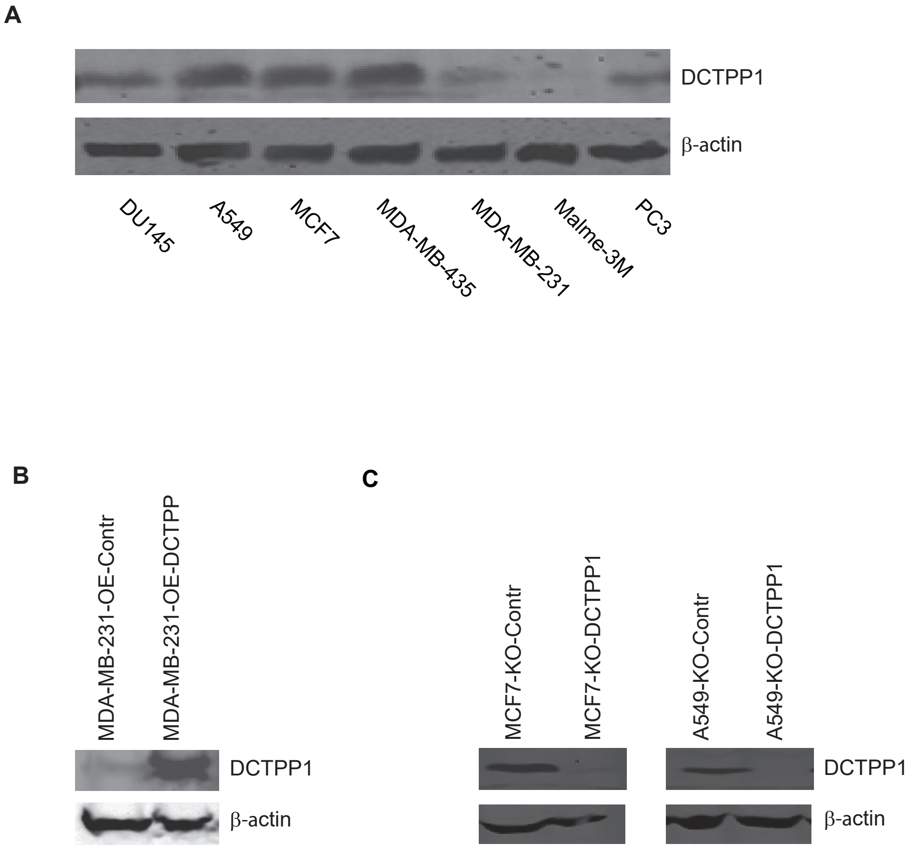


**Figure S12. (Related to Figure 8.) The endogenous DCTPP1 expression level varies among cell lines.** (A). Immunoblotting detection of endogenous expression level of DCTPP1 among a series of cancer cell lines. β-Actin was included as a loading control. (B).Immunoblotting confirmation of lentiviral overexpression of DCTPP1 in MDA-MB-231 cells. β-Actin was included as a loading control. (C). Immunoblotting confirmation of CRISPR-Cas9 knockout of DCTPP1 in A549 and MCF7 cells.

A

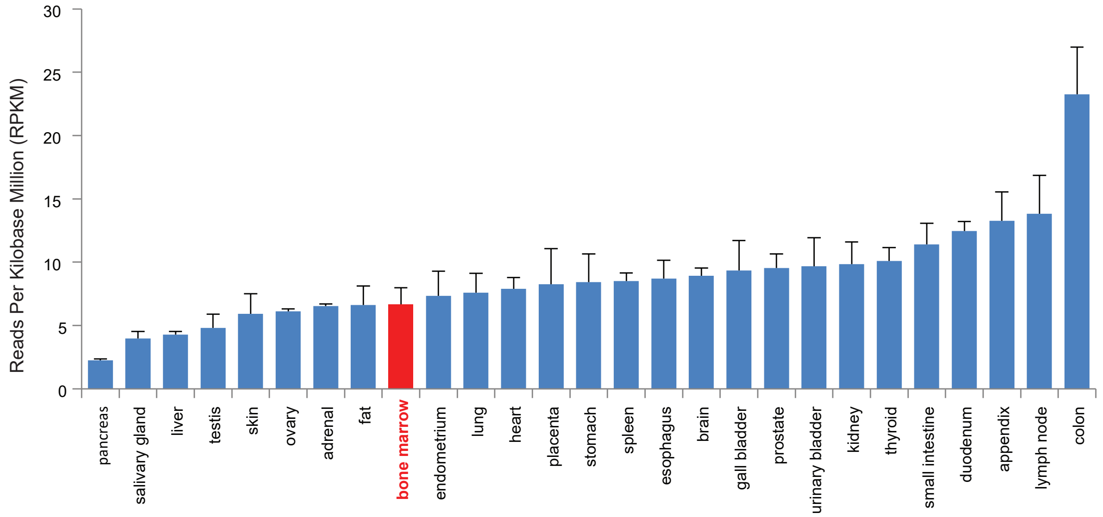


B

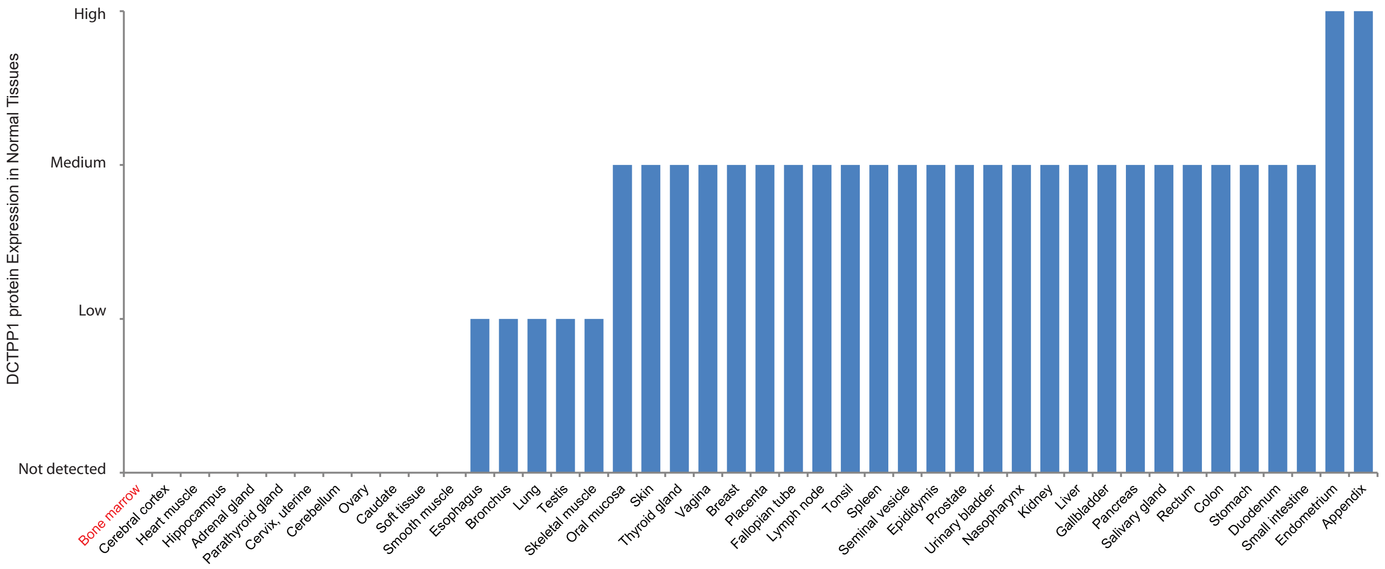


**Figure S13. (Related to Figure 8). DCTPP1 expression levels in hematological tissues are significantly lower than those in most normal human tissues.** (A). DCTPP1 expression levels in bone marrow (Red) are relatively low among most normal human tissues. Human Protein Atlas (HPA) RNA-seq data were downloaded from www.ncbi.nlm.nih.gov/gene/, project: PRJEB4337. Each value represents the mean ± SEM of RNA-seq of multiple human normal tissue samples. (B). DCTPP1 protein expression levels in normal tissues were largely consistent with RNA expression data. DCTPP1 protein expression levels in normal tissues were developed with HPA002832 antibody and were downloaded from www.proteinatlas.org.The protein expression in bone marrow was not detected.

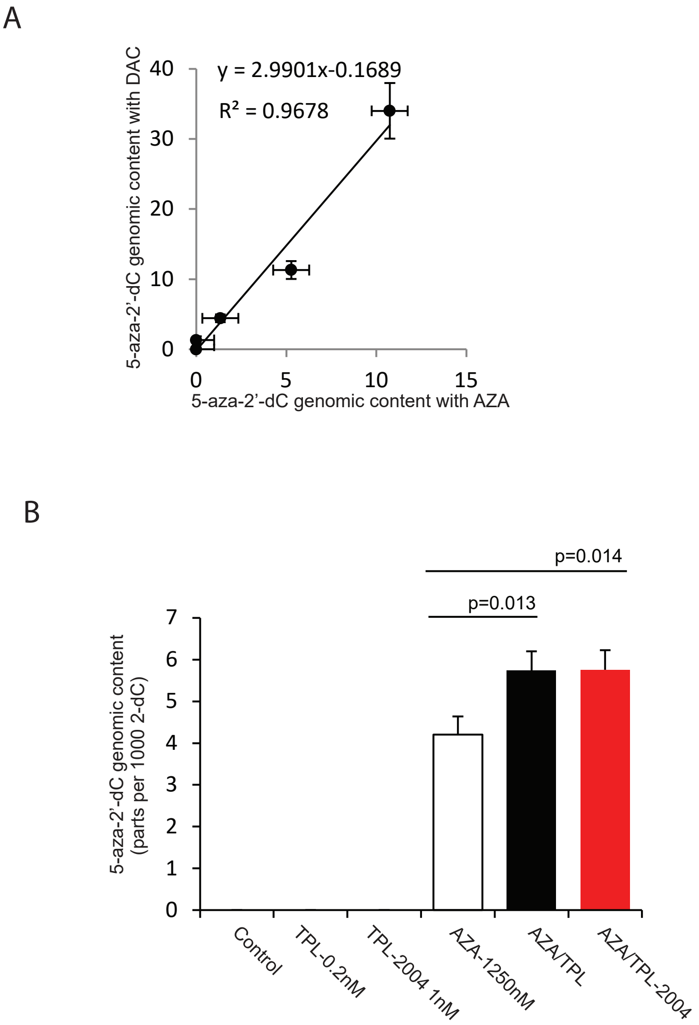


**Figure S14. (Related to Figure 2,4,8.) Combination treatment of azacitidine and triptolide or its DCTPP1 active derivative (TPL-2004) enhanced 5-aza-2'-dC incorporation into genomic DNA.** (A).The potency for 5-aza-2'-dC incorporation into genomic DNA between decitabine *versus* azacitidine was highly correlated .Human prostate cancer DU145 cells were treated with dose series of decitabine or azacitidine for 48 hours, and the 5-aza-2'-dC content in genomic DNA was analyzed with LC-MS/MS. The potency of 5-aza-2'-dC incorporation into genomic DNA across the tested doses of decitabine and azacitidine was highly correlated ($R^2$=0.968), with a ~3-fold greater potency, on average, for decitabine compared to azacitidine. Each point on the graph represents the mean ± SEM of the triplicate measurements for 5-aza-2'-dC content in genomic DNA. (B). 5-aza-2'-dC incorporation into genomic DNA, determined by LC-MS/MS, in DU-145 cells treated with vehicle control, triptolide or its DCTPP1 targeting derivative (TPL-2004), azacitidine alone, or in combination for 2 days. The mean of the amount of 5-aza-2'-dC per 1000 2'-dC in genomic DNA ± SEM of triplicate treatments are shown. Both Triptolide and TPL-2004 treatment significantly enhanced the azacitidine incorporation into genomic DNA in DU145 cells.

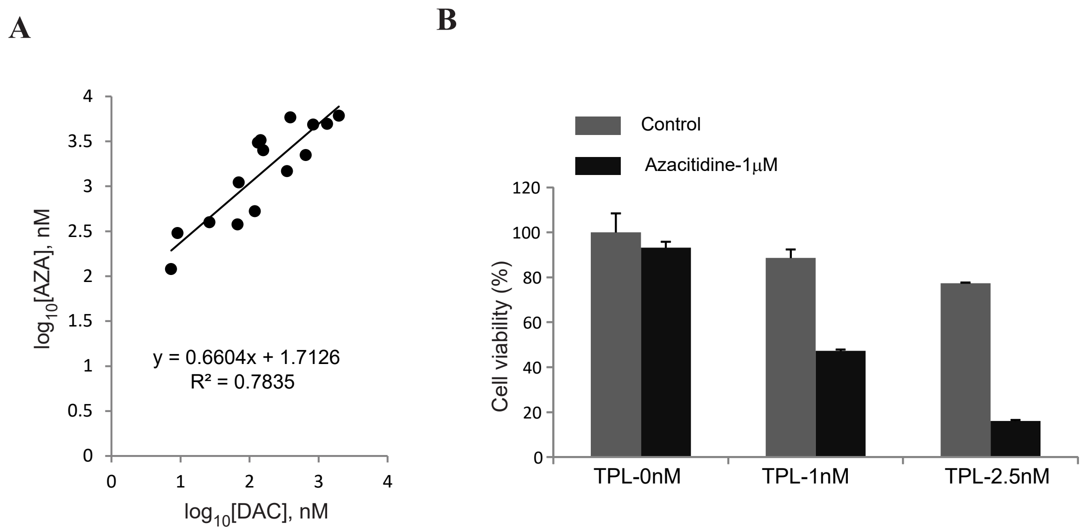


**Figure S15. (Related to Figure 8.) Triptolide synergistically sensitizes cancer cells to nucleoside analog DNMT inhibitor azacitidine *in vitro*.** (A). The potency for growth inhibition between decitabine (DAC) and azacitidine(AZA) was highly correlated. For a panel of 15 cancer cell lines representing multiple cancer types, we measured the potency of decitabine and azacitidine for inducing growth inhibition measured by tritiated thymidine incorporation assay across a broad dose range. The potency of decitabine for growth inhibition (represented as $\log_{10}$ of the $IC_{50}$ concentration in nM) on the panel of cancer cell lines was highly correlated ($R^2$=0.7835, $p < 0.01$) with the potency of azacitidine, with a ~10-fold greater potency, on average, for decitabine compared to azacitidine. Each point on the graph represents the $IC_{50}$ for decitabine *versus* azacitidine for a single cell line. (B). Synergy of azacitidine and triptolide in DU145 cells. Viability measurements of cells treated with vehicle control, 1μM azacitidine or triptolide (1nM and 2.5nM) alone, or in combinations, with each measurement representing the percent viability with respect to the vehicle control treatment. Each value represents the mean ± SEM of triplicate treatments.

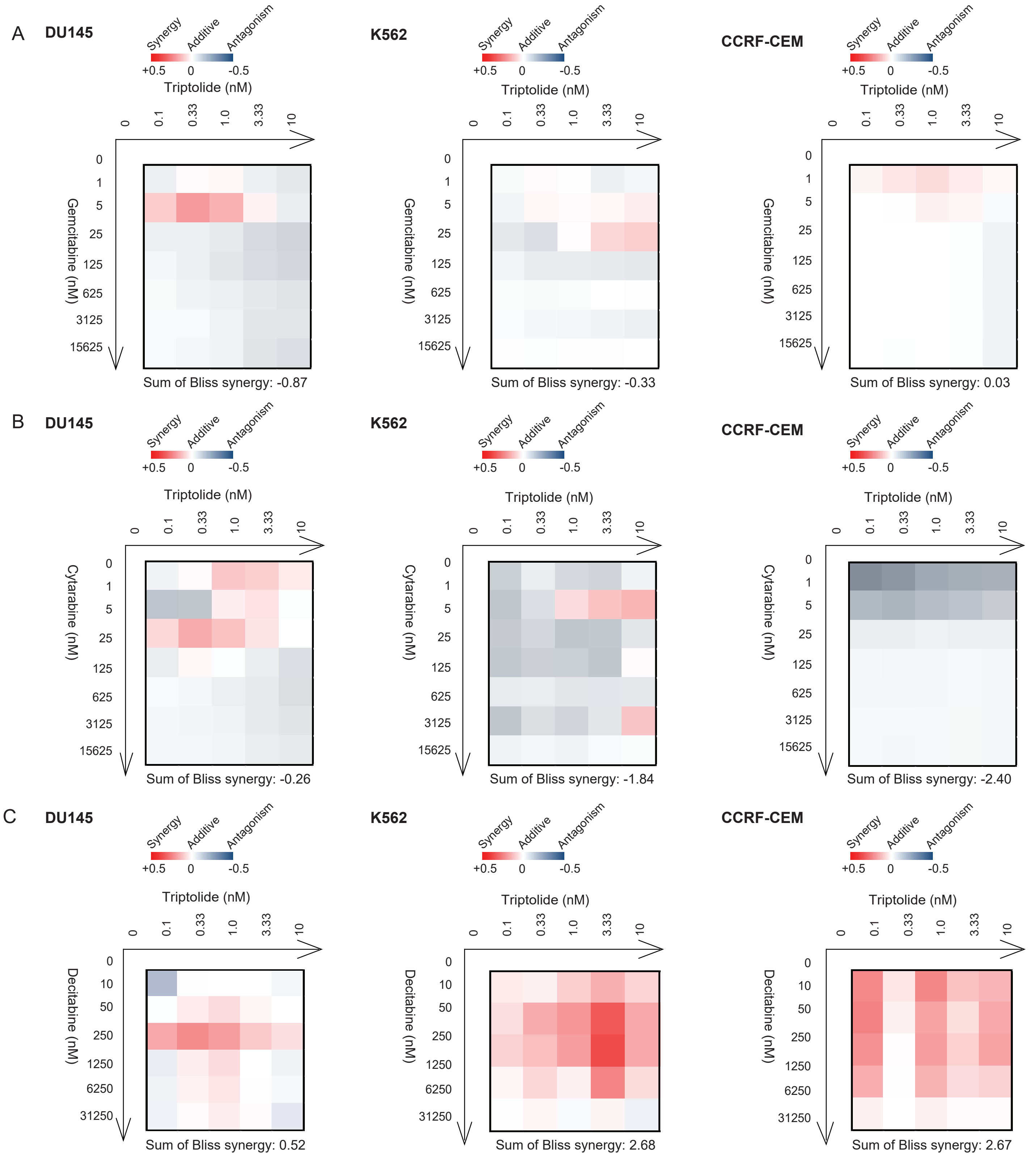


**Figure S16. (Related to Figure 8.) Triptolide does not synergistically inhibit cancer cell growth with gemcitabine and cytarabine.** (A). Bliss synergy analysis of triptolide and gemcitabine (A) or cytarabine (B) in different cancer cell lines. Cell viability measurements of human prostate cancer DU145 cells (left panel), human leukemia K562 cells (middle panel) and CCRFCEM cells (right panel) treated with gemcitabine or cytarabine or triptolide alone and in combinations across a dose series were developed by Alamar Blue Assay. The degree of synergy of combinations of triptolide and gemcitabine (A) or cytarabine (B) was calculated as the Bliss synergy score across the full dose ranges as represented in a heatmap scaled as shown in the color legend. For all 3 cell lines, there was no overall synergy (sum of bliss synergy score <0) for the triptolide combined with gemcitabine or cytarabine. (C). In contrast, there was overall synergy (sum of bliss synergy score >0) between triptolide and decitabine in DU145 cells (left panel), K562 cells (middle panel) and CCRFCEM cells (right panel). Conventions for display are the same as in panels (A) and (B).

| Data collection | DCTPP1$^{(21-130)}$•triptolide |
|---|---|
| Wavelength | 0.92009 |
| Resolution range – Å (last shell) | 29.25 - 2.09 (2.16 - 2.09) |
| Space group | P 1 |
| Monomers/AU | 8 |
| Unit cell (x, y, z – Å; α, β, γ – ˚) | 58.16, 60.92, 73.46; 91.46, 101.31, 104.13 |
| Total reflections | 196314 (17778) |
| Unique reflections | 55230 (5223) |
| Multiplicity | 3.6 (3.4) |
| Completeness – % | 97.6 (92.3) |
| Mean I/σ | 6.73 (1.35) |
| Wilson B-factor | 38.04 |
| R-merge | 0.112 (0.684) |
| R-meas | 0.1321 (0.81) |
| R-pim | 0.0694 (0.429) |
| CC1/2 | 0.989 (0.732) |
| CC* | 0.997 (0.919) |
| Reflections used in refinement | 55210 (5219) |
| Reflections used for $R_{free}$ | 1999 (189) |
| Refinement | |
| $R_{work}$ | 0.187 (0.297) |
| $R_{free}$ | 0.241 (0.363) |
| $CC_{work}$ | 0.961 (0.759) |
| $CC_{free}$ | 0.935 (0.688) |
| *Number of non-hydrogen atoms* | 7225 |
| macromolecules | 6758 |
| ligands | 213 |
| solvent | 254 |
| Protein residues | 855 |
| $RMS_{bonds}$ – Å | 0.009 |
| $RMS_{angles}$ – ˚ | 1.68 |
| Ramachandran favored – % | 97.59 |
| Ramachandran allowed – % | 2.05 |
| Ramachandran outliers – % | 0.36 |
| Rotamer outliers – % | 0.15 |
| Clashscore | 8.8 |
| *Average B-factor* | 42.96 |
| macromolecules | 42.49 |
| ligands | 51.73 |
| solvent | 48.23 |

Supplementary Table 1. Data collection and model refinement statistics. Numbers in parentheses refer to values in the highest resolution shell.
